# Omega-seq: ultra-low-background RNA sequencing with faithful molecular counting and precise transcript-end capture

**DOI:** 10.64898/2026.03.12.711386

**Authors:** Hui-Min Chen, Jui-Chun Kao, Ching-Po Yang, Christabel Tan, Tzumin Lee, Ken Sugino

## Abstract

PCR-based RNA sequencing methods generate phantom unique molecular identifiers (UMIs) when residual UMI-bearing oligonucleotides reprime during amplification, inflating molecule counts and corrupting relative abundances. They also incur non-specific amplification that limits sensitivity, while synthetic poly-T tracts disrupt sequencing and obscure transcript 3′ ends. Here we present Omega-seq, which restricts UMI incorporation to reverse transcription using uracil-containing, USER-excisable template-switching oligonucleotides and a uracil-intolerant polymerase. Omega-seq produces near-background-free libraries, distinguishes 10 fg of input RNA from negative controls, and eliminates bead cleanup. An Omega-shaped dT primer prevents poly-T-induced dephasing, enabling stranded, nucleotide-resolution mapping of transcript ends. Dilution series, ERCC spike-ins, and a rarefaction metric show that Omega-seq has the lowest UMI inflation among methods tested. In individual *Drosophila* neuroblasts, phantom-UMI inflation remains minimal; pooled neuroblast and neuron coverage enables *de novo* discovery of novel multi-exon genes, 3′ UTR extensions, and candidate enhancer RNAs.

---

Single-cell RNA sequencing relies on amplification to recover the minute RNA content of individual cells and on unique molecular identifiers (UMIs) to infer molecular counts [1, 2]. Full-length, plate-based methods commonly use template switching [3] and were initially developed for sensitivity and isoform resolution without molecular counting [4, 5]. UMIs were later incorporated into full-length protocols such as Smart-seq3 [6] and FLASH-seq [7], after becoming standard in droplet-based platforms including 10x Genomics, Drop-seq, and inDrop [8–10]. In each case the UMI is carried by an RT-stage oligonucleotide, either the template-switching oligonucleotide (TSO) or the oligo-dT capture primer, so accurate counting assumes that UMIs are incorporated exclusively during reverse transcription.

Residual UMI-bearing RT oligonucleotides, however, can reprime during downstream PCR, assigning new barcodes to genuine cDNA molecules and generating “phantom UMIs” [6]. This inflates molecule counts, and because generation is stochastic, it also distorts relative abundances and differential-expression estimates. Its mechanism, prevalence, downstream consequences, and limited computational correctability are examined in the accompanying study [11]. Existing mitigation strategies, including competitive inhibition by high primer concentrations and exonuclease digestion, rely solely on reaction kinetics or incomplete enzymatic removal [6, 12], and standard snapshot saturation measures are confounded by sequencing depth and RT efficiency.

Two additional limitations arise from library architecture. Non-specific amplification limits sensitivity at low input and necessitates lossy cleanup before tagmentation. Separately, sequencing through the synthetic poly-T tract of conventional oligo-dT primers causes signal dephasing on Illumina platforms, reducing read quality and obscuring transcription end sites (TES), which limits direct analysis of alternative polyadenylation and 3′ UTR dynamics.

We therefore developed Omega-seq, combining two architectural innovations with one analytical metric (Fig. 1). The Omega-dT primer relocates the sequencing handle to bypass synthetic poly-T, enabling stranded TES mapping. A multi-lock system couples USER-mediated depletion of uracil-containing TSOs, a uracil-intolerant polymerase, and excess PCR primer, restricting productive UMI incorporation to reverse transcription. The logQ-slope summarizes UMI rarefaction kinetics to assess residual phantom-UMI inflation.

**Fig. 1.**
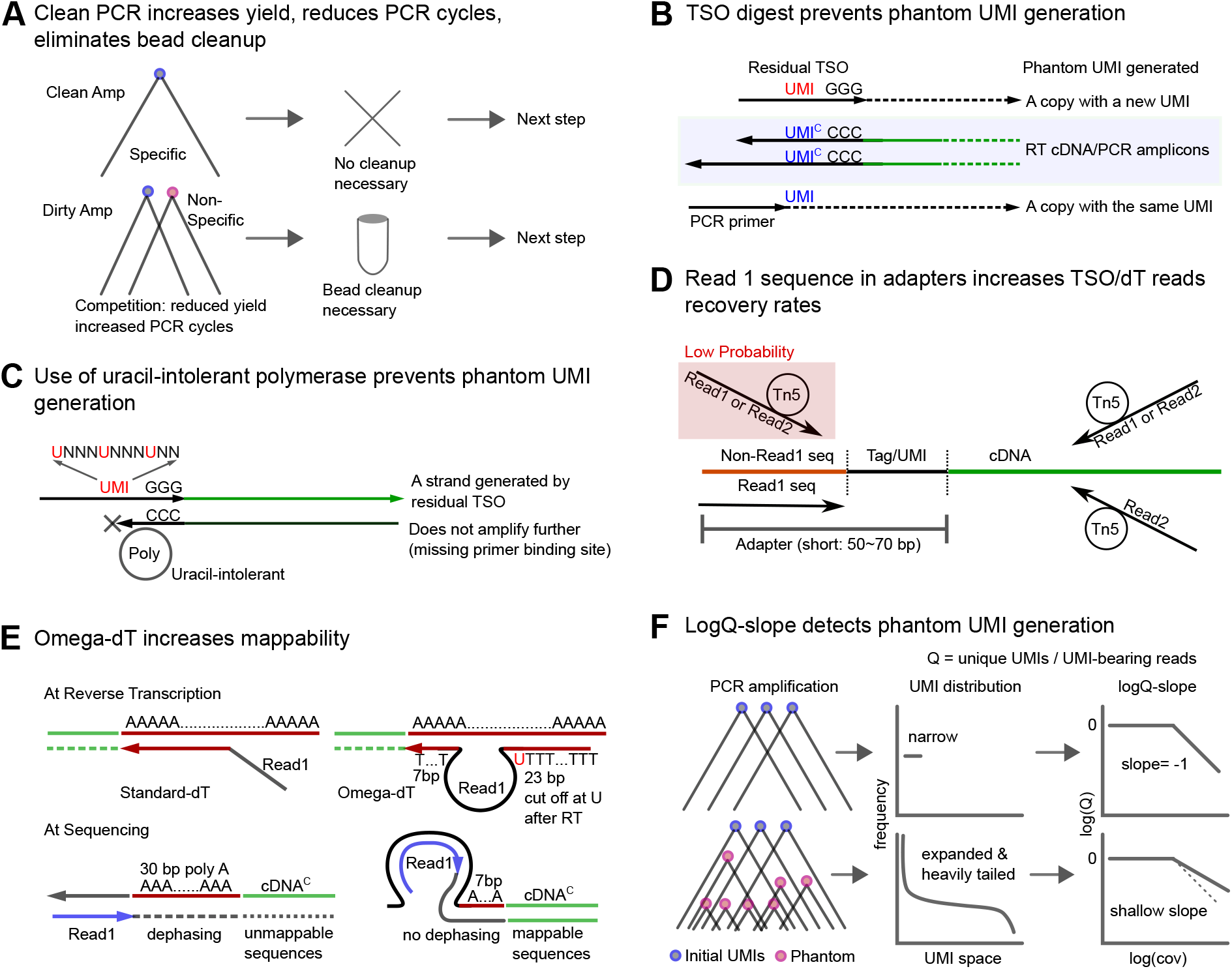
Conceptual framework and architectural innovations of Omega-seq. **A**: Schematic of PCR resource dynamics. In protocols prone to non-specific amplification, stoichiometric competition for reagents (primers, dNTPs, and polymerase) reduces target yield, necessitating compensatory PCR cycles and intermediate purification steps. Omega-seq preserves resource availability by suppressing non-specific products. **B**: Mechanism of TSO-mediated phantom UMI generation. Both the forward (FWD) PCR primer (bottom) and residual TSO (top) can anneal to two types of substrate (shaded): RT cDNAs carrying the TSO complement at their 3′ ends, and PCR amplicons carrying the FWD primer complement at their 3′ ends. High sequence homology between the TSO and 3′ end (e.g., the GGG+spacer) allows residual TSO to sometimes outcompete the FWD primer for priming at these sites, even when longer, higher-affinity FWD primers or elevated primer concentrations are used. Extension from a residual TSO introduces an artifactual “phantom” barcode. **C**: The Omega-seq multi-lock system. The TSO incorporates deoxyuridine (dU) at multiple internal positions (Supp. Fig. 2), including the spacer and UMI regions. This modification, paired with a uracil-intolerant DNA polymerase, provides an additional barrier to amplification from residual TSOs. **D**: Optimization of TSO/dT read recovery. In architectures lacking an integrated Read1 sequence (top), identifiable TSO or dT reads require Tn5 tagmentation within an extremely narrow window (red line), a low-probability event that yields poor recovery of end fragments. Omega-seq incorporates the Read1 sequence directly into the TSO and dT adapters (bottom), removing the requirement for such events and substantially increasing the number of identifiable TSO/dT reads. **E**: The Omega-dT architecture for precise TES mapping. Traditional oligo-dT designs place the sequencing adapter upstream of the homopolymer tract, leading to signal dephasing and unmappable sequences. The Omega-dT primer relocates the adapter handle toward the 3′ terminus, bypassing the poly-A stretch to enable precise, stranded Transcription End Site (TES) determination. **F**: Rarefaction kinetics as a diagnostic of UMI fidelity. In high-fidelity libraries, UMI diversity is established during reverse transcription; subsequent clone-size variation reflects PCR and sequencing stochasticity, producing a relatively narrow distribution. Conversely, phantom UMI generation during PCR creates a heavy-tailed distribution of low-copy, late-cycle barcodes. This divergence is quantitatively captured by the logQ-slope, the slope of the log(*Q*) versus log(coverage) curve, providing a sensitive metric for library purity.

Using ERCC spike-ins, dilution series, and single-cell data, we show that Omega-seq minimizes detectable UMI inflation, produces near-background-free libraries, and enables precise, stranded transcript-end capture. Pooled coverage supports reference-free *de novo* gene-model reconstruction and the discovery of novel genes, unannotated 3′ UTR extensions, antisense transcripts, and candidate enhancer RNAs.

## Results

### Omega-seq suppresses non-specific amplification by oligonucleotide redesign and depletion

Non-specific products generated by primer dimerization, mis-priming, or primer-independent DNA synthesis compete with low-abundance cDNA for finite PCR reagents [13, 14]. Such products may arise during PCR despite hot-start polymerases, or during reverse transcription through the DNA-dependent DNA polymerase activity of the reverse transcriptase [15].

To compare background amplification across current methods, we monitored the amplification of serially diluted total RNA (1000–0.1 pg) and no-template controls in Smart-seq2, Smart-seq3, FLASH-seq-LA, and FLASH-seq-UMI, using SYBR Green real-time PCR and microfluidic capillary electrophoresis. Because the aim was to isolate the contribution of the RT and PCR oligonucleotides themselves, all methods were run in a common buffer, thermal profile, and oligonucleotide concentration (0.3 *µ*M TSO and dT, 1 *µ*M total PCR primer; Methods) rather than at their individually published concentrations. Total primer molarity was matched so that single- and two-primer architectures could be compared at equal oligonucleotide load. This comparison therefore reports the specificity of each oligonucleotide set under matched conditions, not the background of each protocol as published.

All four protocols generated non-specific products, with no-template controls frequently reaching cycle thresholds similar to those of low-input samples (Fig. 2A, upper panels). Capillary electrophoresis revealed broad smears or discrete products predominantly between 50 and 500 bp (Fig. 2A, lower panels).

**Fig. 2.**
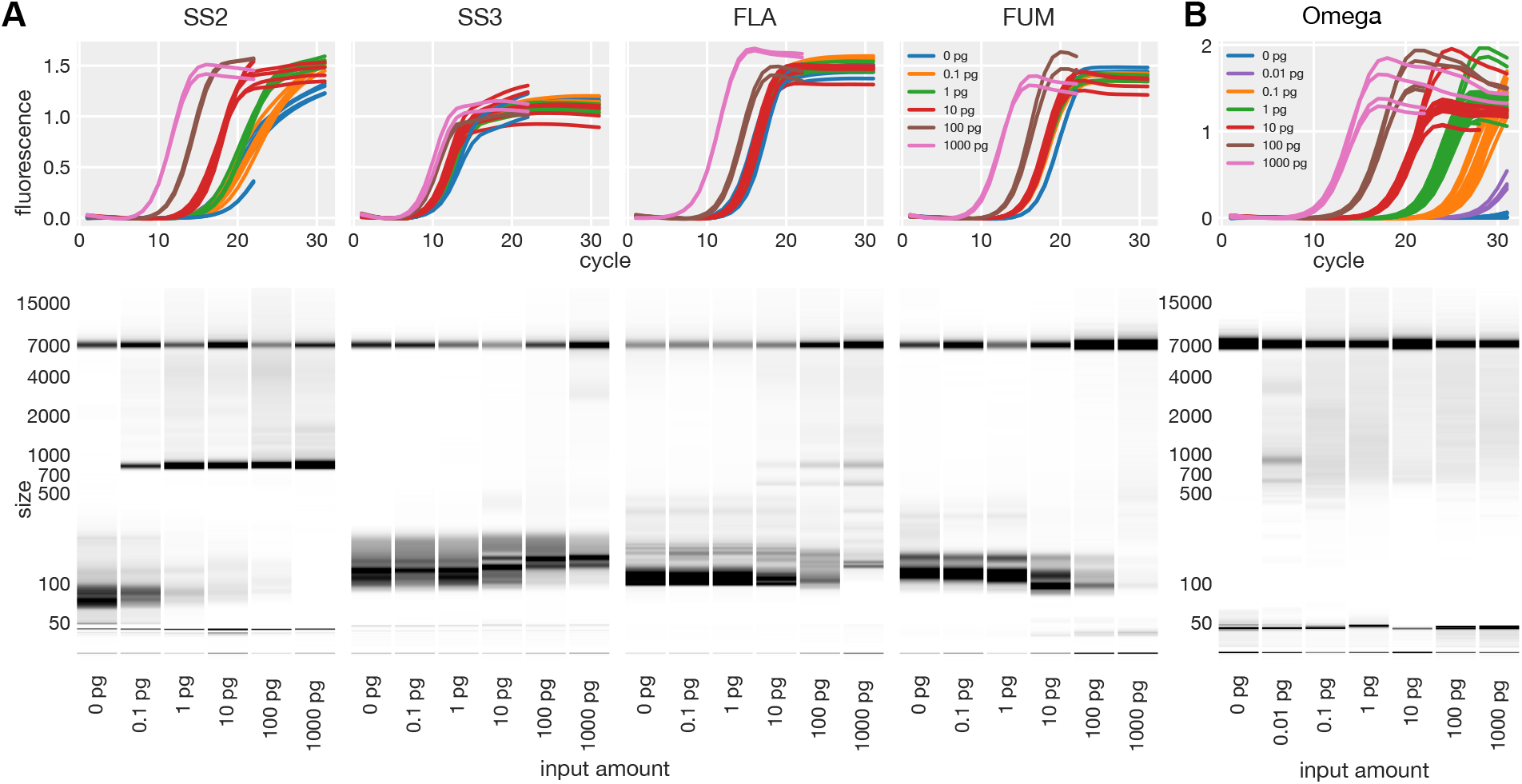
**A**: Oligonucleotide-dependent background across methods. Upper panels: Real-time PCR amplification traces of serially diluted total RNA (1000, 100, 10, 1, 0.1 pg) from *Drosophila* larvae and a negative control (0 pg). SS2: Smart-seq2; SS3: Smart-seq3; FLA: FLASH-seq-LA; FUM: FLASH-seq-UMI. Multiple runs with possibly different total cycles are superimposed. Lower panels: Representative LabChip electropherograms visualized as pseudo-gels. Total area is normalized. Lowest bands and bands at 7000 bp are LabChip markers. Residual primers appear near 50 bp. **B**: Suppression of oligonucleotide-derived background in Omega-seq. Real-time PCR traces (upper) and LabChip profiles (lower) demonstrating clean separation between 0.01 pg input and negative controls. (Note: in addition to the standard set of inputs ranging from 0.1 pg to 1000 pg, 0.01 pg was added.) Importantly, the uracil-intolerant polymerase acts only in the full Omega-seq protocol; the qPCR comparisons in panels A and B used a uracil-tolerant polymerase (Methods), so they report oligonucleotide specificity rather than this amplification lock, whose contribution is assessed within the full protocol (Fig. 4).

Single-primer PCR methods (Smart-seq2, FLASH-seq-LA) showed lower background than two-primer systems, consistent with “pan-handle suppression,” in which complementary adapter ends form a stem-loop that inhibits primer annealing and extension [16], and with fewer opportunities for off-target interaction when only one primer sequence is present. Random UMI sequences in Smart-seq3 and FLASH-seq-UMI may further increase non-specific hybridization.

We therefore designed Omega-seq around a single-primer PCR architecture combined with enzymatic depletion of residual RT oligonucleotides. Iterative optimization (Supp. Figs. 1, 2; Supp. Note 2) produced amplification curves that scaled with input concentration while negative controls remained flat (Fig. 2B), distinguishing 10 fg of input RNA from negative controls by both qPCR and electrophoresis.

Three design features were central. First, the Illumina s5 sequence was fused to the Tn5 Mosaic Element to create a single ISPCR handle and improve recovery of TSO- and dT-containing fragments (Fig. 1D). Second, selected thymines in the TSO and dT primers, including positions within the UMI (UNNNUNNNUNN), were replaced with uracil and cleaved with USER enzyme [17]. Thermolabile exonuclease I, as used in SCRB-seq [12], instead reduced gene-detection sensitivity, possibly through degradation of cDNA ends or ISPCR primers despite phosphorothioate protection [18]. Third, replacing the variable `VN` oligo-dT anchor with a fixed `TT` terminus suppressed RT-derived products formed in the absence of RNA [15, 19].

Uracil-based depletion was previously used in Smart-seq-total [20], but there the UMI is carried by the oligo-dT primer and becomes covalently incorporated into first-strand cDNA, so uracil excision would cleave the UMI-labelled cDNA itself; residual UMI-bearing oligo-dT primers therefore persist and can generate 3′ phantom UMIs during PCR. Omega-seq instead places the UMI on the TSO, whose sequence is copied into cDNA while the oligonucleotide itself remains separate, so residual UMI-bearing TSOs can be neutralized by USER excision and uracil-intolerant amplification. Because Omega-dT carries no UMI, residual oligo-dT primer cannot create new molecular identities.

### Nanopore characterization of preamplified cDNA

The preceding qPCR and electrophoresis results describe the amplification reaction rather than the sequenced library. To examine the preamplified product directly, we sequenced it on the Oxford Nanopore platform, omitting the intermediate purification, tagmentation, and index PCR steps that would otherwise alter its composition. We compared Omega-seq with FLASH-seq-UMI, the closest current-generation full-length protocol carrying UMIs, using serially diluted total RNA (10, 1, and 0.1 pg) and no-template controls, 20 PCR cycles for both, and pooled loading on a single flow cell. This characterizes the material entering library preparation and is not a comparison of final Illumina libraries: in routine use, FLASH-seq-UMI includes an SPRI bead cleanup at this stage that removes short products, and the profiles below therefore describe the material that cleanup is required to remove.

Approximately 4 million reads had the expected adapter orientation and structure (Fig. 3B, Supp. Note 3) and were evenly distributed across samples (Fig. 3A, Supp. Fig. 3A), except the Omega-seq 0 pg control, which yielded only ∼ 2,000 reads despite equivalent loading. Its barcode–adapter crossover rate was more than 100-fold higher than in other conditions (Fig. 3D), suggesting that these reads largely arose from index hopping rather than true amplification, consistent with the near-flat negative controls observed by qPCR.

**Fig. 3.**
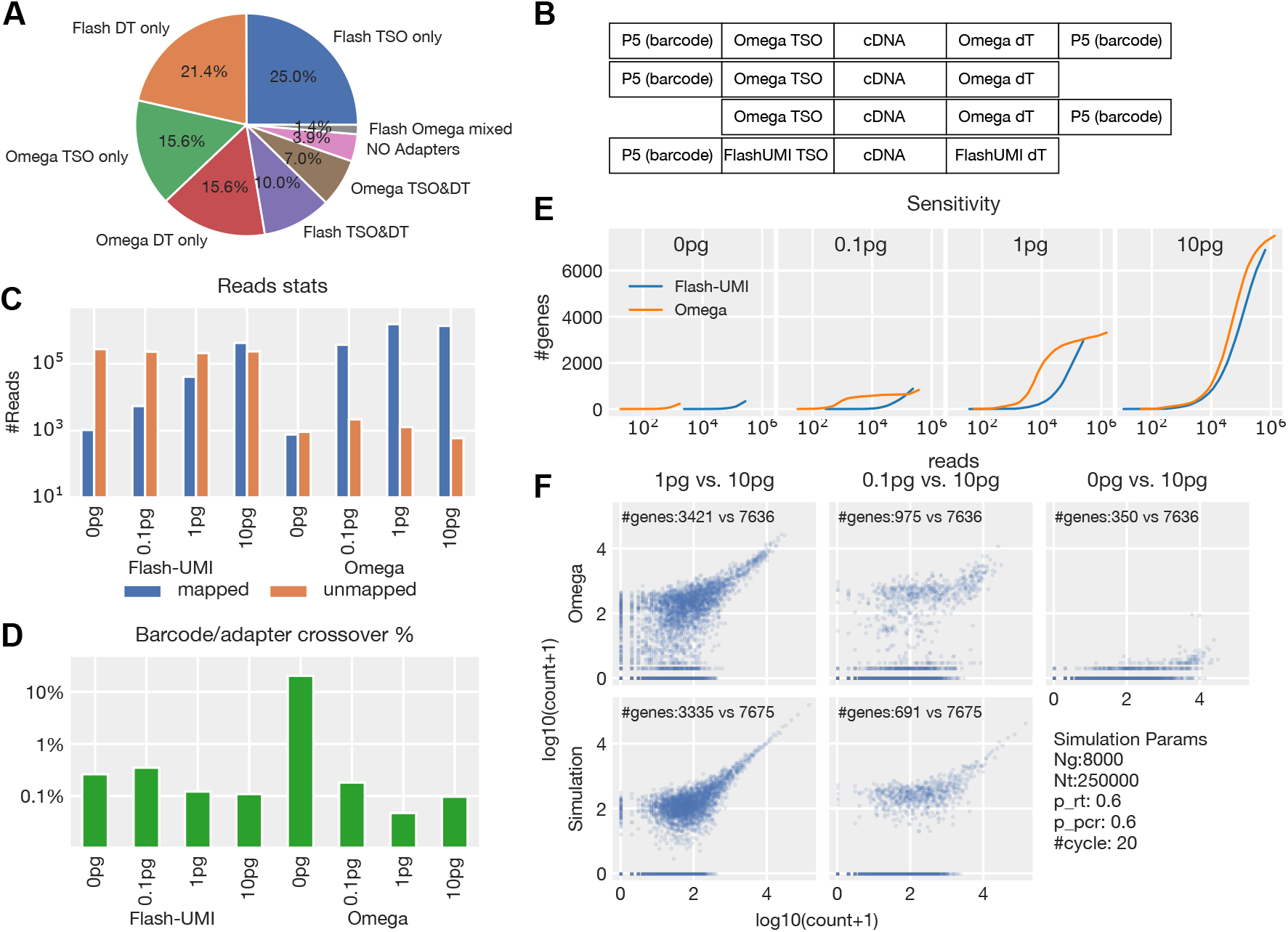
Nanopore characterization of preamplified cDNA. **A**: Adapter composition of total Nanopore reads, classified by detected adapter types (FLASH TSO, FLASH dT, Omega TSO, Omega-dT) and their respective combinations. **B**: Structural read filtering. Only reads possessing the proper adapter ordering and orientation are extracted for downstream analysis (maximum 1 bp gap allowed between detection units). **C**: Total number of mapped (blue) and unmapped (orange) reads plotted on a log scale, highlighting the collapse of mappability in low-input FLASH-seq-UMI preamplified products. **D**: Barcode adapter inconsistency rate, quantifying cross-contamination (e.g., a FLASH-seq-UMI sample inappropriately containing an Omega-seq adapter sequence). **E**: Gene detection rarefaction curves. The number of detected genes is plotted against sequencing depth for FLASH-seq-UMI and Omega-seq across a serial dilution (10, 1, and 0.1 pg) and no-template control (0 pg) of *Drosophila* total RNA. **F**: (Upper panels) Scatter plots of log-transformed gene counts comparing the 10 pg input (x-axis) against lower inputs and negative controls, illustrating the stochastic sampling plateau, clearly visible in the middle panel. (Lower panels) Corresponding scatter plots generated from simulated data, accurately recapitulating the experimental stochastic bottleneck.

In unpurified preamplified material, more than 99% of Omega-seq reads mapped to the genome at all non-zero input concentrations (Fig. 3C, Supp. Fig. 3B), whereas FLASH-seq-UMI mappability fell from 65% at 10 pg to 16% at 1 pg and 2% at 0.1 pg. The unmapped FLASH-seq-UMI reads consisted predominantly of inserts shorter than 50 bp, including primer dimers and sequences matching bacterial or vector contaminants. Omega-seq also maintained high-molecular-weight insert distributions characteristic of full-length cDNA (Supp. Fig. 3C, D), matching the capillary electrophoresis profiles (Fig. 2B).

Gene-detection rarefaction curves computed from these unpurified products showed earlier saturation for Omega-seq at 1 pg and 0.1 pg (Fig. 3E). Restricting the FLASH-seq-UMI analysis to inserts longer than 500 bp, which approximates the effect of the SPRI cleanup its protocol specifies, restored comparable sensitivity (Supp. Fig. 4A). The difference therefore reflects the burden of short artifacts rather than reduced cDNA recovery; purification removes this burden in FLASH-seq-UMI, whereas it is largely absent from Omega-seq.

**Fig. 4.**
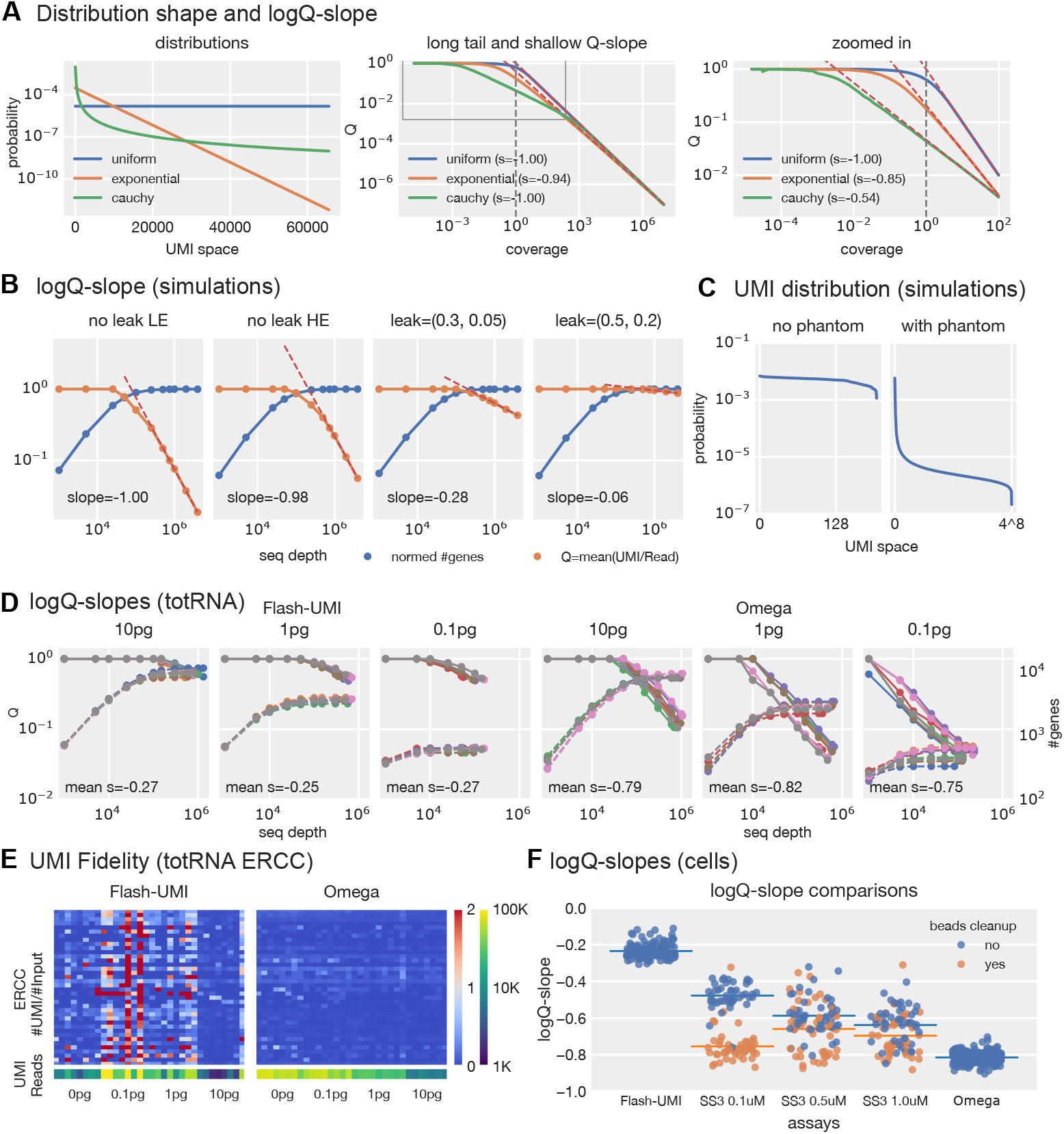
Quantification of UMI fidelity via logQ-slope analysis. **A**: Theoretical logQ-slope behavior. For various distributions with differing heavy-tailedness (leftmost panel), Q vs. coverage (=sequence reads / initial copy number) is plotted (middle and right panels). Note that while all distributions eventually reach a slope of − 1 in log-log space, the gradient of the initial decay phase is highly dependent on the heavy-tailedness of the distribution. The vertical dashed line indicates the “standard saturation point” (coverage = 1). **B**: LogQ-slope analysis for simulated data with parameters: num genes=8000, num transcripts=250,000. **Panel 1:** No phantom UMI generation, low RT efficiency (p_rt=0.3), low PCR efficiency (p_pcr=0.7). **Panel 2:** No phantom UMIs, high efficiencies (p_rt=0.9, p_pcr=0.9). **Panel 3:** High efficiencies with moderate phantom UMI generation (w_tso=0.3, w_pcr=0.05). **Panel 4:** High efficiencies with severe phantom UMI generation (w_tso=0.5, w_pcr=0.2). Orange lines represent Q; blue lines show the number of genes detected. (For parameter details, see Supp. Note 5) **C**: Simulated UMI distributions for the most abundant gene corresponding to Panel 1 (left) and Panel 3 (right) in (B). Note the difference in x range. **D**: LogQ-slope analysis for 10/1/0.1 pg human total RNA sequenced on the Illumina platform (n = 8 each). Solid lines represent Q (left y-axis); dashed lines show genes detected (right y-axis). Slope values between FLASH-seq-UMI and Omega-seq are significantly different (*p <* 2.5 × 10^−7^). **E**: Absolute fidelity of ERCC UMIs (Observed UMIs / Input copy number) displayed as a heatmap (y-axis representing ERCC species and x-axis representing samples). Red shading indicates values *>*1, with values larger than 2 capped as solid red (indicating severe phantom UMI inflation). Samples consist of serially diluted human total RNA and negative controls spiked with 0.05 *µ*l of a 1:72,000 ERCC dilution. **F**: LogQ-slopes for single-cell data. FLASH-seq-UMI and Omega-seq data were generated from individual *Drosophila* neuroblasts (sample sizes indicated in Methods). The middle three columns represent Smart-seq3 data [6] separated by PCR primer concentrations. Yellow dots indicate samples subjected to pre-PCR bead cleanup. Note: FLASH-seq-UMI parameters reflect their current protocol (0.5 *µ*M forward and 0.1 *µ*M reverse PCR primer, 1.84 *µ*M TSO) compared to Omega-seq (1 *µ*M ISPCR primer, 0.3 *µ*M TSO).

Across the dilution series, gene-count scatter plots showed a horizontal plateau, most clearly in the 10 pg versus 0.1 pg comparison (Fig. 3F, upper panels), as expected when dilution reduces transcripts spanning a broad abundance range to single molecules before PCR. Simulations reproduced the plateau closely (Fig. 3F, lower panels; Supp. Fig. 4B), supporting a stochastic sampling limit rather than an amplification artifact; a small number of points below it may reflect inefficient amplification, late-cycle chimeras, or index hopping. Omega-seq therefore recovers transcripts present as single molecules at the onset of PCR.

### Rarefaction and ERCC analyses demonstrate high UMI fidelity in Omega-seq

Smart-seq3 suppresses phantom-UMI generation through competition from high concentrations of PCR primer [6, 21]. Omega-seq retains this competition (1 *µ*M ISPCR primer) and adds two further biochemical barriers: USER digestion of the uracil-containing TSOs, and amplification with a uracil-intolerant polymerase (KAPA HiFi, standard formulation), which together block amplification of residual oligonucleotides.

To assess UMI fidelity, we developed the logQ-slope, the log–log slope of *Q* = *U/R* against sequencing depth, where *U* is the number of unique UMIs and *R* the number of UMI-bearing reads. When UMIs are assigned before PCR, discovery saturates with increasing depth and the slope approaches − 1. Phantom UMIs generated throughout PCR instead form a heavy-tailed distribution of low-copy barcodes that continue to be discovered, producing a shallower slope (Fig. 4A–C). Simulations across leak rates, transcript copy numbers, and PCR efficiencies show that the slope responds far more strongly to repriming on primer-ended amplicons (*w*_pcr_) than on TSO-ended amplicons (*w*_tso_), and that its behaviour is stable over a wide parameter range, degrading only at very low PCR efficiency, very low copy number, or when UMI collisions saturate the barcode space (Supp. Fig. 5). The underlying branching process, coverage dependence, and relationship to clone-size distributions are described in the accompanying study [11].

**Fig. 5.**
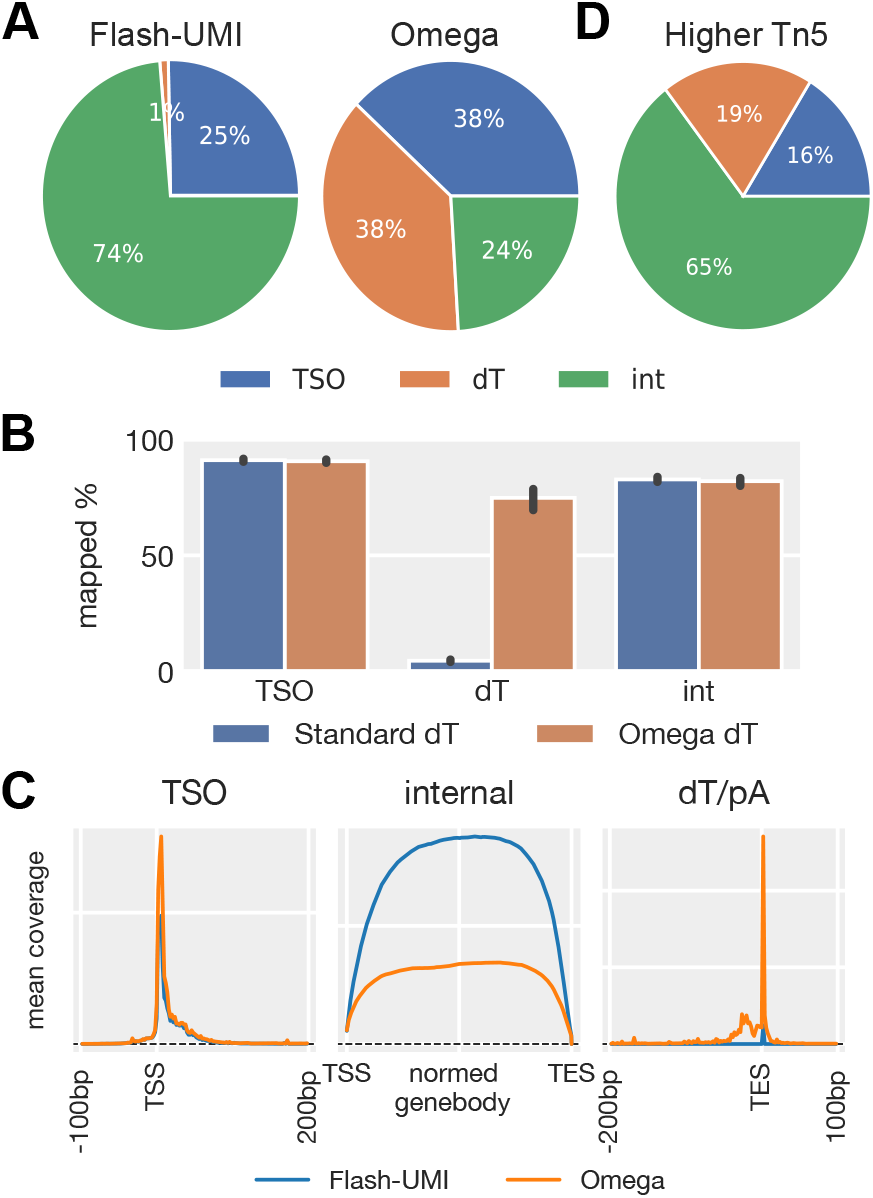
**A**: Sequence reads distribution. Proportion of reads assigned to TSO-derived (containing 5′-tag), dT-derived (containing 3′-adapter or *>*20 bp poly-T), or internal fragments (lacking terminal tags). Data for panels A–D were generated using high-depth Illumina sequencing. **B**: Impact of primer architecture on genomic mappability. Comparison of mapping rates for TSO, internal, and dT-derived reads. Note the failure of standard dT primers due to Illumina dephasing, contrasted with the high mappability recovered by the Omega-dT structural design. **C**: End-to-end coverage profiles. Normalized read density mapped across gene bodies for TSO, internal, and dT-derived fragments. Peak densities sharply align at annotated TSS and TES boundaries, demonstrating successful stranded, dual-end capture. **D**: Impact of Tn5 concentration on the proportion of recovered TSO/dT end-reads. The lower Tn5 concentration condition corresponds to the data shown in (A).

We evaluated the metric using high-depth Illumina libraries generated from serially diluted human total RNA (10, 1, and 0.1 pg) and no-template controls (*n* = 8 per condition), each containing ERCC spike-ins. FLASH-seq-UMI, run at its published oligonucleotide concentrations (0.5 *µ*M forward PCR primer, 1.84 *µ*M TSO), produced shallow logQ-slopes of approximately − 0.3, whereas Omega-seq slopes approached − 0.8 (Fig. 4D).

ERCC spike-ins provided an independent absolute benchmark. We calculated the fidelity ratio as detected UMIs divided by input ERCC molecules (Fig. 4E). Many FLASH-seq-UMI transcripts exceeded a ratio of 1, in some cases reaching 172, demonstrating extensive molecular overcounting. Omega-seq ratios remained below 1 across the tested abundance range, consistent with restriction of UMI incorporation primarily to reverse transcription.

The ERCC ratio is conservative: because the spike-in is added at a fixed amount while input varies, ERCC receives a smaller share of reads at higher inputs, and low RT efficiency or shallow ERCC sampling can then keep the ratio below 1 even when phantom UMIs are present, as in the 10-pg FLASH-seq-UMI samples. The logQ-slope instead reads the shape of the rarefaction curve and remains informative once coverage passes the saturation bending point. It is used here as a *relative*, coverage-matched comparison between libraries prepared and sequenced side by side, not as an absolute score against a fixed cutoff: the clean baseline itself drifts from near − 1 at deep coverage towards ≈ − 0.7 when reads per UMI are few, so a single global threshold misclassifies a large fraction of shallow but clean libraries (Supp. Fig. S3 of the companion study [11]). The rarefaction series is therefore evaluated over a common set of sequencing depths for every library, and slopes are compared only between libraries prepared and sequenced together.

Omega-seq and FLASH-seq-UMI also differ roughly 12-fold in the primer-to-TSO ratio that governs competitive suppression (0.27 versus 3.3), so the logQ-slope difference in Fig. 4D could reflect oligonucleotide concentrations alone. We therefore used published Smart-seq3 datasets [6] to separate primer competition from the biochemical barriers, alongside FLASH-seq-UMI and Omega-seq libraries from individual *Drosophila* neuroblasts (Fig. 4F). Smart-seq3 slopes steepened progressively with increasing PCR-primer concentration, confirming the competitive axis. Pre-PCR SPRI bead cleanup steepened them further, but slopes varied widely between cleaned samples at the same primer concentration, indicating that physical TSO removal is technically sensitive and not uniformly complete. Omega-seq values were tightly distributed and steeper than both the Smart-seq3 1-*µ*M primer condition (*p <* 10^−80^) and the cleanest bead-cleaned Smart-seq3 condition (0.1 *µ*M forward primer; *p <* 10^−18^), indicating that the multi-lock strategy suppresses post-RT UMI generation beyond what primer competition or physical cleanup achieves. An independent estimator agrees: applying the clone-size detector of the companion study [11] to these same libraries scores Omega-seq among the cleanest of the 23 datasets it surveys (*ŵ*_tso_ ≈ 0.05, versus ≈ 0.4 for FLASH-seq-UMI and for Smart-seq3 at 0.1 *µ*M forward primer; its Fig. 2A), and the per-library logQ-slopes and *ŵ*_tso_ values rank the shared libraries almost identically (Spearman *ρ* = 0.90; its Supp. Table B3).

### Omega-dT enables stranded capture of transcription end sites

Smart-seq3 and FLASH-seq-UMI enrich 5′-end reads by incorporating Illumina s5 and Tn5 Mosaic Element (ME) sequences into the TSO and forward PCR primer, but direct capture of transcript 3′ ends remains difficult. Appending the s5+ME sequence to the 5′ end of a conventional oligo-dT primer increased recovery of dT-derived reads, but these reads mapped poorly, because sequencing through the synthetic poly-T tract caused signal dephasing and loss of base quality [22, 23] (Fig. 5A,B; Supp. Fig. 2).

We therefore repositioned the adapter internally within the oligo-dT primer, creating an omega-shaped loop upon hybridization to the poly-A tail (Fig. 1E; Supp. Figs. 1, 2). This geometry places the sequencing start site adjacent to the transcript boundary and bypasses the synthetic poly-T tract. Omega-dT consequently restored the mappability of dT-derived reads and produced sharp, stranded coverage peaks at annotated TES positions, complementing TSO-derived TSS reads to provide end-to-end transcript capture (Fig. 5B,C). The slightly lower mappability of Omega-dT reads relative to TSO and internal reads likely reflects occasional sequencing into the endogenous poly-A tail on the opposite strand. Consistent with genuine 3′-end capture, Omega-dT read positions clustered at annotated TES and at the upstream AATAAA polyadenylation signal (Supp. Fig. 6A,B). A separate population mapped to genomic A-rich tracts (Supp. Fig. 6C), reflecting internal priming; these reads are removed by the local sequence-context filter described in Methods.

**Fig. 6.**
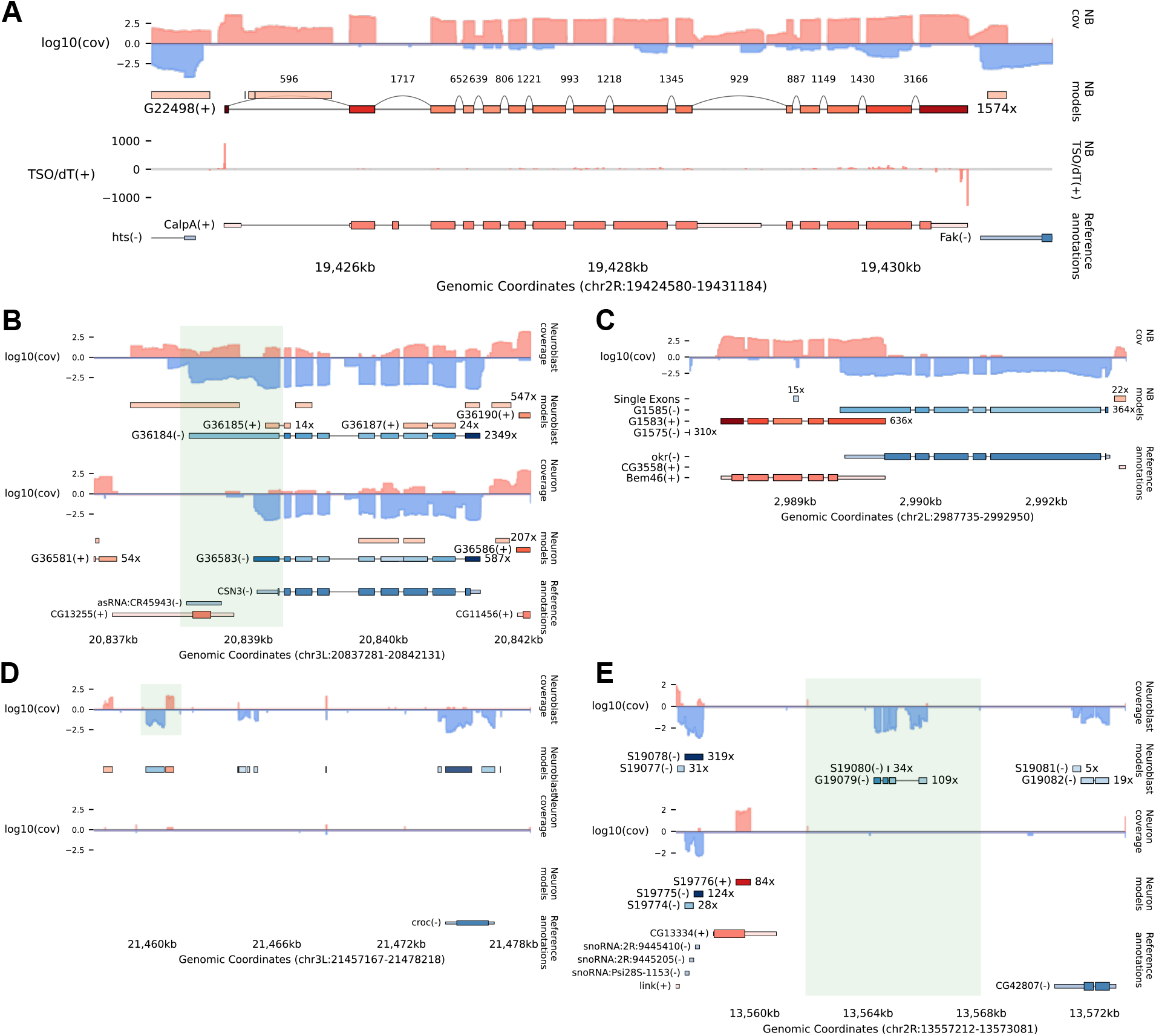
**A**: Representative example of *de novo* gene model reconstruction. In addition to internal splice junctions, TSO and dT read positions are integrated to accurately define the absolute boundaries of 5′ and 3′ exons. TSO and dT read mapping positions are displayed in the “TSO/dT(+)” track, with TSO counts plotted as positive values and dT counts as negative values for visual clarity. Only collapsed models are shown for both the reconstructed and reference gene tracks. In general, data mapping to the positive (+) strand is indicated with red shading, while data mapping to the negative (-) strand is indicated with blue shading. **B**: Identification of a novel, elongated 3′ UTR. Note that this specific 3′ UTR extension is differentially expressed between *Drosophila* neuroblasts and neurons. **C**: Accurate resolution of physically overlapping transcripts on opposing strands. **D**: Identification of a candidate eRNA, observable exclusively in the neuroblast population. **E**: Discovery of a novel, unannotated multi-exon gene detected specifically in neuroblasts.

The balance between internal and terminal reads could be tuned through Tn5 concentration. Internal fragments require both s5- and s7-loaded transposition events, whereas terminal fragments already contain an s5 handle and require only s7 transposition (Fig. 1D). Higher Tn5 concentrations favored internal fragments, whereas lower concentrations enriched TSS- and TES-containing reads (Fig. 5A,D).

### Stranded, end-delineated reads enable *de novo* gene reconstruction

We developed a custom *de novo* reconstruction pipeline that integrates pooled TSO- and Omega-dT-defined transcript boundaries, splice junctions, and stranded read coverage across cells. Terminal reads define 5′ and 3′ exons, internal exons are delimited by splice junctions, and the resulting elements are assembled into gene models through a strand-specific splice graph. Terminal reads were first filtered by local sequence context to exclude priming artifacts, in particular dT reads at genomic A-rich tracts (Methods).

Pooling reads across *Drosophila* neuroblasts and neurons provided sufficient coverage for reconstruction. TSO and Omega-dT reads also impart strand orientation to their internal paired-end mates, extending stranded coverage across gene bodies. This enabled accurate delineation of transcript boundaries in reconstructed models (Fig. 6A; Supp. Fig. 7) and revealed features not represented in standard gene-by-cell matrices or reference annotations: differentially expressed 3′ UTR extensions (Fig. 6B; Supp. Fig. 8), overlapping transcripts resolved on opposite strands (Fig. 6C; Supp. Fig. 9), neuroblast-specific candidate enhancer RNAs (Fig. 6D; Supp. Fig. 10), novel multi-exon genes differentially expressed between the two populations (Fig. 6E; Supp. Fig. 12), and unannotated antisense transcripts within introns of known genes (Supp. Fig. 11).

## Discussion

PCR-based single-cell RNA sequencing has two distinct architectural vulnerabilities. Residual UMI-bearing oligonucleotides can reprime during PCR, assigning new molecular identities to existing cDNA molecules and distorting relative abundances. Separately, conventional oligo-dT designs force sequencing through synthetic poly-T tracts, causing dephasing and obscuring transcript 3′ ends. Omega-seq addresses both through coordinated redesign of the reverse-transcription oligonucleotides and the amplification scheme, while also suppressing non-specific amplification.

The low background of Omega-seq has both analytical and practical consequences. Suppressing non-specific products increases the fraction of informative reads and avoids the losses associated with purification, especially at very low RNA inputs. Direct progression from PCR to tagmentation, which removes a purification step required by protocols that generate short products, also simplifies automation and reduces reagent use and manual handling.

Omega-dT addresses a separate limitation of conventional oligo-dT priming. By positioning the sequencing handle at an internal position proximal to the primer’s 3′ terminus, it bypasses the synthetic poly-T tract and enables high-quality sequencing across the transcript–poly-A junction. Together with TSO-derived 5′ reads, Omega-dT provides stranded information at both transcript termini and imparts strand orientation to internal paired-end reads. Pooling this coverage across cells enabled reference-free *de novo* gene-model reconstruction and the identification of novel multi-exon genes, 3′ UTR extensions, antisense transcripts, and candidate enhancer RNAs in *Drosophila* neurons and neuroblasts. Omega-seq therefore combines molecular counting with the structural information required to define features absent from existing annotations.

Because phantom-UMI generation is stochastic, it distorts relative abundances and fold-changes rather than simply scaling total counts; its mechanism, prevalence, downstream consequences, and computational detection are examined in the accompanying study [11]. Omega-seq addresses the artifact experimentally through USER digestion of uracil-containing TSOs, amplification with a uracil-intolerant polymerase, and competitive inhibition by excess PCR primer. Together these barriers restrict productive UMI incorporation to reverse transcription and also remove a promiscuous primer that can generate non-specific products and spurious transcript-start signals.

We use the logQ-slope as a rarefaction-based summary of UMI fidelity. Unlike snapshot saturation measures such as *Q* = *U/R*, it captures the continued discovery of low-abundance UMIs produced by phantom generation. Its theoretical basis, coverage dependence, and relationship to full clone-size-distribution modelling are described in the accompanying study [11]. Here, together with ERCC spike-ins, it provides an orthogonal benchmark showing that Omega-seq minimizes detectable post-RT UMI inflation.

These findings extend to droplet-based platforms, where the accompanying study detects a reproducible residual-oligonucleotide phantom signature in current 10x GEM-X 3′ data [11]. Because UMI counts underlie gene-by-cell matrices, the artifact can affect relative expression, differential analysis, and measurements of cellular heterogeneity.

The two principal Omega-seq innovations transfer differently to 3′ and 5′ droplet assays. Omega-dT geometry could be incorporated into bead-bound 3′ capture primers to bypass synthetic poly-T and enable direct TES sequencing, but it would not eliminate phantom UMIs. As in Smart-seq-total, the barcode- and UMI-bearing oligo-dT primer is covalently incorporated into cDNA, so unused primer cannot be selectively destroyed without also damaging the corresponding primer-derived sequence in productive cDNA molecules. In 3′ droplet assays, the additional tethering of the primer to the gel bead ensures that residual unused primer is carried into the amplification reaction. In 5′ assays, the UMI is instead carried on a TSO, the configuration exploited by Omega-seq, and the multi-lock system should therefore be directly adaptable in principle. Whether a UMI is copied or incorporated thus determines which strategy applies.

Several limitations remain. Omega-seq was developed in a plate-based, template-switching setting, and its performance across other cell types, oligonucleotide concentrations, and amplification chemistries remains to be established. The multi-lock system also constrains polymerase choice to uracil-intolerant high-fidelity enzymes. TES calls remain subject to internal priming at genomic A-rich tracts (Supp. Fig. 6C), which we filter computationally rather than prevent. The logQ-slope requires coverage beyond the saturation bending point and reports a relative signature rather than a calibrated phantom fraction, and its dynamic range narrows when the barcode space approaches saturation or amplification is inefficient (Supp. Fig. 5). Finally, *de novo* gene models depend on pooled rather than per-cell coverage, and the novel features they reveal are candidates pending independent confirmation.

The workflow could also be made more scalable. Cellular barcodes could be inserted into the TSO spacer and the Omega-dT loop, permitting pooling immediately after reverse transcription and reducing the number of separate PCR reactions. Because such barcodes identify the cell rather than the molecule, they do not compromise UMI fidelity even if residual primer survives. This strategy is particularly suited to tagmentation with an s7-only transposome: only fragments containing either the TSO or Omega-dT end provide the complementary s5/Read 1 handle required to generate P5 during index PCR. Every retained read pair therefore carries strand information together with a cellular barcode. A fully RNA-based TSO (Supp. Fig. 2, V9-TSO) could also provide a lower-cost alternative to USER digestion, since residual RNA TSOs can be removed with RNase I.

Overall, Omega-seq shows that persistent limitations of PCR-based RNA sequencing can arise from oligonucleotide architecture rather than amplification efficiency alone. Controlling the fate of UMI-bearing primers improves molecular-counting fidelity, while repositioning the sequencing handle enables direct transcript-end capture. These principles extend beyond the present protocol: Omega-dT could add TES information to 3′ droplet assays, whereas the multi-lock TSO strategy is more directly transferable to 5′ assays.

## Supplementary information

Supplementary information is available for this paper.

## Declarations

- Funding: HHMI, NIH R01(NS134890)
- Conflict of interest/Competing interests: K.S. and T.L. have filed a provisional patent related to the Omega-dT primer.
- Ethics approval and consent to participate. Not applicable. This study used *Drosophila melanogaster*, which is not subject to institutional animal care and use regulation, and commercially sourced, deidentified human total RNA (Takara Bio #636530). No human subjects research was performed and no IRB approval was required.
- Consent for publication. Not applicable.
- Data availability: All sequencing data have been deposited to GEO under accession number GSE[XXX].
- Materials availability. Oligonucleotide sequences sufficient to reproduce Omega-seq are provided in Supp. Fig. 2. A provisional patent application covering the Omega-dT primer has been filed; oligonucleotides and protocol materials are available from the corresponding author for non-commercial research use under a standard materials transfer agreement.
- Code availability: All analysis code and logQ-slope pipeline are available on GitHub at [XXX].
- Author contributions: K.S. designed the oligonucleotides and the Omega-seq protocol, conceived and supervised the experiments, analyzed the data, and wrote the manuscript. H.C. and J.K. performed the experiments and generated the sequencing libraries. C.Y. and C.T. performed cell sorting. T.L. secured funding, contributed to intellectual discussions, and supervised the overall progress of the project.

## Online Methods

### Sample preparation

Total RNA was extracted from *Drosophila* larval brains using the Quick-DNA/RNA Microprep Plus kit (Zymo Research, #D7005) and quantified using a Qubit fluorometer (Thermo Fisher Scientific).

*Drosophila* larvae of genotype UAS-mCD8::GFP, ase-Gal4/CyO [24] were hatched within a 2 h window and reared at 25^°^C for 50 h. Brains from 50 h after larval hatching (ALH) were dissected, the ventral nerve cord was removed, and the tissue was digested and triturated according to previously published methods [25]. The dissociated cells were then FACS sorted into 384-well plates using a BD FACSDiscover S8 Cell Sorter (BD Biosciences).

### qPCR

Real-time PCRs to assess oligo performance for various methods were performed using the same buffer composition. Buffer, cycling, and oligonucleotide concentrations were held constant across methods by design, so that differences in background reflect oligonucleotide architecture rather than protocol-specific reaction conditions. Published method-specific concentrations were used only where stated (Fig. 4F). The RT reaction mix contained: 5% PEG8000, 0.075% Triton X-100, 25 mM Tris-HCl (pH8.3), 30 mM NaCl, 2.5 mM MgCl_2_, 1 mM GTP, 8 mM DTT, 0.5 U/*µ*l RNase Inhibitor, 0.5 mM each of dNTPs, and 2 U/*µ*l Maxima H Minus reverse transcriptase. Oligonucleotide concentrations were 0.3 *µ*M each for TSO and dT. The RT thermal profile was 42^°^C 90 min; 10 cycles of (50^°^C 2 min, 42^°^C for 2 min); 85^°^C for 5 min; and a 4^°^C hold. This RT buffer mostly follows the Smart-seq3xpress composition [21].

The qPCR buffer consisted of 0.02 U/*µ*l Phanta High-Fidelity DNA polymerase (uracil-tolerant), 1 × Phanta buffer, 1 × SYBR Green, and 0.2 mM each of dNTP. Because only the Omega-seq oligonucleotides carry uracil, 0.05 U/*µ*l USER enzyme (NEB) was supplemented into the Omega-seq reaction alone; all other components, cycling, and oligonucleotide concentrations were identical across methods. The total PCR primer concentration was 1 *µ*M in all reactions: 1 *µ*M of the single ISPCR primer for single-primer architectures (Smart-seq2, FLASH-seq-LA, Omega-seq), and 0.5 *µ*M each of forward and reverse primer for two-primer architectures (Smart-seq3, FLASH-seq-UMI). The PCR thermal profile was 37^°^C for 1 hour (only if using USER enzyme); 95^°^C for 2 min; followed by 22 to 32 cycles of (95^°^C for 15 sec, 65^°^C for 20 sec, 72^°^C for 4 min). End products were diluted, and their size distributions were characterized using a LabChip GX Touch II (Revvity).

### Nanopore sequencing

Two replicates of each serially diluted total RNA input (10 pg, 1 pg, and 0.1 pg) from *Drosophila* larval CNS and negative controls (0 pg) were reverse transcribed and PCR amplified according to FLASH-seq-UMI and Omega-seq protocols, both utilizing 20 cycles for PCR. Following initial amplification, 10 *µ*l (2 × 5 *µ*l) of FLASH-seq-UMI product was re-amplified using 0.5 *µ*M each of barcoded-P5 and DI-PCR-P1A-R (FLASH-seq-UMI REV primer) for 16 cycles in a 30-*µ*l KAPA HiFi PCR reaction. For Omega-seq, 8.2 *µ*l (2×4.1 *µ*l) of the amplified product was re-amplified using 1 *µ*M barcoded-P5 primer for 16 cycles in a 30-*µ*l KAPA HiFi PCR reaction.

The samples within each method were pooled, cleaned using the DNA Clean & Concentrator-5 kit (Zymo Research), and quantified using Qubit. Subsequently, 100 ng of DNA from each method was combined as input for Nanopore Ligation Sequencing Kit V14 (Oxford Nanopore Technologies). The resulting library was sequenced using Nanopore MinION for 72 hours.

### Illumina sequencing

Human brain total RNA (Takara Bio, #636530) was serially diluted to final amounts of 10 pg, 1 pg, and 0.1 pg and added to the lysis mix in place of water. ERCC spike-ins (0.05 *µ*l of a 1:72,000 dilution) were included in the RT mix for both protocols. Eight replicates of each dilution and eight negative controls were processed by each method, using a Mantis liquid dispenser (Formulatrix), a Mosquito liquid handler (SPT Labtech), a Bio-Rad S1000 Thermal Cycler, and a QuantStudio 5 PCR system (Thermo Fisher Scientific). Omega-seq libraries were prepared as described in Supp. Note 1, with preamplification products proceeding directly to Tn5 tagmentation and index PCR without bead cleanup. FLASH-seq-UMI samples were processed at the published buffer and primer concentrations and were subjected to SPRIselect bead cleanup after preamplification PCR and before tagmentation. Indexed libraries were quantified using Qubit, their size distributions characterized on a LabChip GX Touch II, and submitted to GENEWIZ for Illumina sequencing. Sorted *Drosophila* neuroblasts and neurons were processed similarly.

### Analysis

A custom Python pipeline (GitHub XXX) was developed to detect, classify, and trim various adapter sequences. Adapter-trimmed sequences were mapped using HISAT2 [26] and kallisto [27] for Illumina sequencing, or minimap2 [28] for Nanopore sequencing. To assign mapped reads to genes, we utilized kallisto output coupled with the hg38 or dm6 annotation for Illumina data. For Nanopore data, we developed a custom script utilizing HTSeq [29] to assign reads to transcript isoforms. Custom Python scripts were also used to assign TSO- and dT-derived reads stringently to their respective TSS and TES boundaries.

logQ-slope. For each library, UMI-bearing reads were pseudoaligned to the transcriptome with kallisto and grouped by transcript, and UMIs were error-corrected by directional (Hamming-distance-1) collapse (Supp. Note 4). A rarefaction series was built by randomly subsampling reads without replacement to ten depths (1,000; 5,000; 10,000; 25,000; 50,000; 100,000; 150,000; 300,000; 600,000; and 1,000,000 UMI-bearing reads, each capped at the library size). At each depth we computed, per gene, the total read count and the number of distinct collapsed UMIs, and defined Q as the median across genes (present at more than four depths) of the per-gene ratio of distinct UMIs to reads. The logQ-slope was obtained by fitting ln Q against ln(depth) over every sliding window of three consecutive depths and taking the minimum (steepest) slope. The theoretical basis of this metric is given in the companion study [11].

For *de novo* gene model reconstruction, a custom Python pipeline was developed to aggregate 1) overall read coverage, 2) exact TSO/dT read coordinates, and 3) splice junctions. Briefly, gene models were reconstructed through the following computational steps:

1. **Filtering:** Collected splice junctions were retained only when supported by at least 10 spliced reads and when the spliced-read count was at least 15% of the larger of the two connected exons’ peak coverage; introns shorter than 35 bp were discarded. TSO and dT reads were filtered based on local genomic sequence context to remove priming artifacts (TSO reads for which more than three of the adjacent five bases were G or C, and dT reads with more than five A residues on the positive strand, or five T residues on the negative strand, within the downstream 8 bp, were excluded; this threshold was set from the downstream A-content distribution of dT read positions; Supp. Fig. 6C).
2. **Interval Extraction:** For each chromosome and strand, base intervals where reads formed a contiguous covered range were extracted.
3. **Exon Definition:** Within each contiguous base interval, 5′ exons, 3′ exons, and internal exons were explicitly defined utilizing a) splice junctions, b) TSO positions, and c) dT positions.
4. **Single-Exon Extraction:** In covered regions falling outside of the exons defined in Step 3, single-exon genes were extracted.
5. **Splice Graph Assembly:** Directed splice graphs were constructed between the non-single-exon elements.
6. **Gene Model Generation:** Final gene models were extracted as connected components from the splice graph.

Splice graph path selection (exact isoform determination) was not attempted, as it is inherently imprecise with short-read data; however, the collapsed *de novo* gene models successfully map structural boundaries useful for downstream applications. The Jupyter notebooks detailing the preprocessing steps and figure generation have been deposited in the GitHub repository [XXX].

Statistics. Group comparisons of per-cell logQ-slopes used two-sided, two-sample Student’s t-tests (`scipy.stats.ttest_ind`), treating each cell (or, for the 10 pg total-RNA titration, each replicate library) as an independent observation. Fig. 4D compares n = 8 FLASH-seq-UMI versus n = 8 Omega-seq total-RNA libraries; Fig. 4F compares Omega-seq *Drosophila* neuroblasts (n = 309 cells, one plate) against Smart-seq3 HEK293 cells at 1 *µ*M forward primer (n = 48) and bead-cleaned at 0.1 *µ*M forward primer (n = 48; the cleanest of the no-TSO groups); the FLASH-seq-UMI column in Fig. 4F comprised n = 144 cells.

## Supplementary Notes

### Supplementary Note 1: Omega-seq protocol

(Note: Lysis and RT buffers are based on Smart-seq3xpress [21])

#### Cell Lysis and RNA Denaturation

Prepare the Lysis Mix according to the components listed in Table 1 (1*×* vol in *µ*l).

- **For purified RNA:** Dilute the template RNA in nuclease-free water and add it directly to the prepared Lysis Mix.
- **For single cells:** Sort individual cells directly into PCR tubes or multi-well plates containing the Lysis Mix.

**Table 1.** Lysis Mix (1 × conc=final concentration in the 1.1 *µ*l RT reaction)

| reagent | conc | $1 \times$ conc | $1 \times$ vol ($\mu\text{l}$) |
| --- | --- | --- | --- |
| TritonX-100 | 10% | 0.075 | 0.00825 |
| RNase Inhibitor | 40U/$\mu\text{L}$ | 0.5 | 0.01375 |
| oligo-dT | 10$\mu\text{M}$ | 0.3 | 0.033 |
| H<sub>2</sub>O |  |  | 0.545 |
| Total |  |  | 0.6 |

Incubate the samples at 72^°^C for 3 minutes to facilitate lysis. Proceed immediately to the Reverse Transcription step.

#### Reverse Transcription (RT)

Prepare the RT Master Mix as detailed in Table 2. Add the RT mix to the lysed samples, vortex gently, and spin down. Incubate in a thermal cycler with the heated lid set to 50^°^C using the following program:

**Table 2.** RT Mix

| reagent | conc | 1× conc | 1× vol (μl) |
| --- | --- | --- | --- |
| PEG8000 | 40% | 5 | 0.1375 |
| Tris-HCl(pH8.3) | 1000mM | 25 | 0.0275 |
| NaCl | 1000mM | 30 | 0.033 |
| MgCl<sub>2</sub> | 100mM | 2.5 | 0.0275 |
| GTP | 100mM | 1 | 0.011 |
| DTT | 100mM | 8 | 0.088 |
| dNTPs | 25mM/each | 0.5 | 0.022 |
| MaximaH- | 200U/μL | 2 | 0.011 |
| RNase Inhibitor | 40U/μL | 0.5 | 0.01375 |
| ERCC | 1× | 1/72000 | 0.05 |
| TSO | 10μM | 0.3 | 0.033 |
| H<sub>2</sub>O |  |  | 0.04575 |
| From Lysis |  |  | 0.6 |
| Total |  |  | 1.1 |

1. 42^°^C for 90 min (cDNA synthesis)
2. 10 cycles of:
  - 50^°^C for 2 min
  - 42^°^C for 2 min
3. 85^°^C for 5 min (RT inactivation)
4. Hold at 4^°^C

#### USER Digestion and PCR Amplification

Prepare the PCR Master Mix containing USER enzyme according to Table 3. Add the mix to the RT reaction product. Place samples in a thermal cycler with the heated lid set to 100^°^C. The program below combines the enzymatic cleanup of uracil-containing oligonucleotides with selective amplification:

**Table 3.** PCR Mix

| reagent | conc | 1× conc | 1× vol (μl) |
| --- | --- | --- | --- |
| USER | 1U/μL | 0.05 | 0.205 |
| KAPA HiFi | 2× | 1 | 2.05 |
| ISPCR | 25μM | 1 | 0.164 |
| H<sub>2</sub>O |  |  | 0.581 |
| From RT |  |  | 1.1 |
| Total |  |  | 4.1 |

1. 37^°^C for 60 min (USER digestion)
2. 98^°^C for 3 min (initial denaturation)
3. PCR Amplification (15–20 cycles):
  - 98^°^C for 20 sec (denaturation)
  - 65^°^C for 20 sec (annealing)
  - 72^°^C for 6 min (extension)
4. 72^°^C for 5 min (final extension)
5. Hold at 4^°^C

### Supplementary Note 2: Evolution of Omega-seq oligo design

The development of the final Omega-seq oligonucleotide architecture involved iterative optimization to balance sensitivity, specificity, and PCR efficiency. The sequence logic and rationale for each design iteration are detailed below and sequences and their structures are summarized in Supp. Fig. 2.

#### Nomenclature and Functional Elements

The following abbreviations are used to describe functional sequence motifs:

- **ME**: 19-bp Tn5 Mosaic Element (essential for transposase recognition).
- **s5**: 3′ portion of the Illumina P5 adapter sequence.
- **s7**: 3′ portion of the Illumina P7 adapter sequence.
- **s5+ME**: The compound sequence serving as the Illumina Read 1 sequencing primer binding site.

#### Design iterations and rationale

##### V1 & V2 (Initial s5+ME Integration)

Modified from the standard Smart-seq3 architecture, these designs incorporated the **s5+ME** sequence directly into the TSO and oligo-dT to enable single-primer PCR. *Observation:* The 3′ terminus of the TSO (containing the 8N UMI + GGG) generated a high fraction of reads mapping to non-TSS genomic locations. This indicated significant off-target activity, likely driven by “strand invasion” during reverse transcription or non-specific priming during downstream PCR. **Note:** The 5′-biotin modification prevents TSO concatemerization. Without this modification, continued terminal-cytosine addition and repeated template switching generate TSO concatemers at the 5′ end.

##### V3 & V4 (Spacer Integration)

Inspired by the FLASH-seq-UMI design, we relocated the UMI and inserted a spacer sequence to decrease mis-priming. *Observation:* These constructs caused severe suppression of PCR amplification when using KAPA HiFi polymerase. We hypothesized this was due to strong competitive inhibition from residual oligonucleotides that were not effectively cleared or suppressed. **V5 & V6 (Tm Reduction):** We reduced the total oligonucleotide length to lower the melting temperature (Tm), aiming to decrease the stability of non-specific hybrids and alleviate competitive inhibition during PCR. *Observation:* This modification was insufficient to restore PCR efficiency, suggesting the inhibition was driven by high molar concentrations of residual oligos rather than thermodynamic stability alone. **V7-dT (**3′**-Anchor Optimization):** We modified the 3′-anchoring sequence of the oligo-dT primer from the canonical degenerate `VN` to a fixed `TT` dinucleotide. *Observation:* This change significantly reduced the formation of oligo-dT-derived primer dimers. It suggests that the degenerate VN sequence acts as a promiscuous template for the RT enzyme even in the absence of RNA, whereas the fixed TT anchor suppresses this background activity. **V7-TSO & V8-TSO (Uracil-Mediated Suppression):** To definitively overcome the competitive inhibition observed in V3-V6, we increased the number of uracil bases incorporated into the TSO sequence. *Observation:* Higher uracil content rendered the residual TSOs more susceptible to USER enzyme digestion, successfully restoring PCR efficiency and library yield. **V8-dT (Proof-of-Concept Omega-dT):** The initial design of the Omega-dT primer, featuring an internal adapter sequence to form a loop structure upon poly-A binding. **V9-dT (**3′**-Tag & Poly-T Removal):** An optimized Omega-dT design incorporating a specific 3′-tag to facilitate adapter identification. This version was designed such that the poly-T tail is detached from the amplicon following USER digestion, aiming to minimize possible sources of non-specific products. **V10-dT (All-RNA Poly-T):** A variant where the entire poly-T tract consists of RNA bases to facilitate complete removal during RNase/USER treatment. **V9-TSO (All-RNA TSO):** A fully RNA-based TSO variant designed to test if replacing DNA bases with RNA could further reduce background by enabling RNase-based cleanup (e.g., RNase I).

The final Omega-seq protocol uses V8-TSO and V9-dT.

### Supplementary Note 3: Nanopore structural reads extraction and filtering

Analysis of the raw Nanopore dataset (18.6 million reads) revealed that approximately 80% of reads lacked full-length structural integrity, defined as the concurrent presence of both TSO and oligo-dT adapter sequences (Fig. 3A). Furthermore, an inspection of sample barcodes identified a subset of reads associated with multiple conflicting barcodes or internal adapter sequences. These artifacts typically arise from “chimeric reads” formed during the ligation library preparation step or failures in the base-calling software to correctly segment independent DNA molecules from the raw continuous signal trace [30].

To mitigate these artifacts and isolate high-confidence transcripts, we implemented a strict structural filtering pipeline. We extracted only those sequences containing the expected library topology (Barcode– Adapter–Transcript–PolyA–Adapter), allowing for a maximum tolerance of a 1 bp gap between elements (Fig. 3B). This filtration yielded approximately 4 million structurally complete reads, which were uniformly distributed across sample conditions, with the exception of the Omega-seq 0 pg negative control (Supp. Fig. 3A).

However, even within this structurally filtered subset, we observed instances of discordance between sample barcodes and library adapters (Supp. Fig. 3A). These events are likely attributable to “barcode crosstalk” or index-hopping, driven by sequencing errors within the barcode region that lead to incorrect demultiplexing. This phenomenon has been well-characterized in multiplexed Nanopore sequencing [31]. Notably, the Omega-seq 0 pg condition exhibited a ∼ 100-fold higher rate of barcode/adapter discordance compared to other samples (Fig. 3D). This strongly suggests that the few properly structured reads assigned to the negative control (orange bar, Omega-seq 0 pg) are largely the result of bioinformatic carryover or physical contamination from other multiplexed samples rather than genuine amplification from the blank input.

### Supplementary Note 4: UMI deduplication and error correction

Sequencing error can inflate UMI counts. To correct for this, we performed UMI collapsing using the “directional” adjacency method implemented in UMI-tools [32]. This algorithm corrects for PCR amplification errors by modeling the relationship between UMI sequences as a directed network.

Briefly, reads are first grouped by transcript. Within each group, unique UMIs are identified and their abundance (read count) is tabulated. A directed network is then constructed where nodes represent unique UMIs and edges represent putative error pathways. A directed edge is created from UMI A (source) to UMI B (target) if and only if:

1. The Hamming distance between sequence A and sequence B is exactly 1.
2. The abundance of UMI A (*C*_*A*_) relative to UMI B (*C*_*B*_) satisfies the inequality: *C*_*A*_ ≥ 2*C*_*B*_.

This threshold serves to distinguish true low-copy molecules from PCR or sequencing errors derived from high-copy parents. Finally, all nodes within a connected component are collapsed to the single UMI with the highest read count within that component.

### Supplementary Note 5: Stochastic simulation of library preparation and sequencing

The logQ-slope panels in Fig. 4B were generated with a stochastic simulation of single-cell RNA-seq library preparation that follows the branching-process model of phantom-UMI generation developed in the companion study [11]; we summarize only the parameters here. Transcript abundances are initialized from a Zipf rank–abundance law [33] over *N*_*g*_ genes and *N*_*t*_ total transcripts (`num_genes`, `num_transcripts`). Reverse transcription is a binomial capture with per-molecule efficiency *p*_rt_, and PCR is a Galton–Watson branching process with per-cycle efficiency *p*_pcr_ [34]. Phantom UMIs arise from residual-oligo repriming during PCR, modelled as a two-state (TSO-ended / primer-ended) multi-type branching process. A free TSO reprimes TSO-ended templates at rate *w*_tso_ and PCR-primer-ended templates at rate *w*_pcr_, with each event assigning a new random UMI to a genuine template. The full transition kinetics are derived in the companion study [11]. Sequencing is multinomial sampling of the post-PCR pool to a total depth `seq_depth`. The per-panel values of *p*_rt_, *p*_pcr_, *w*_tso_ and *w*_pcr_ are listed in the Fig. 4 legend. The complete Python implementation is provided on GitHub [XXX].

## Supplementary Figures

**Fig. 1.**
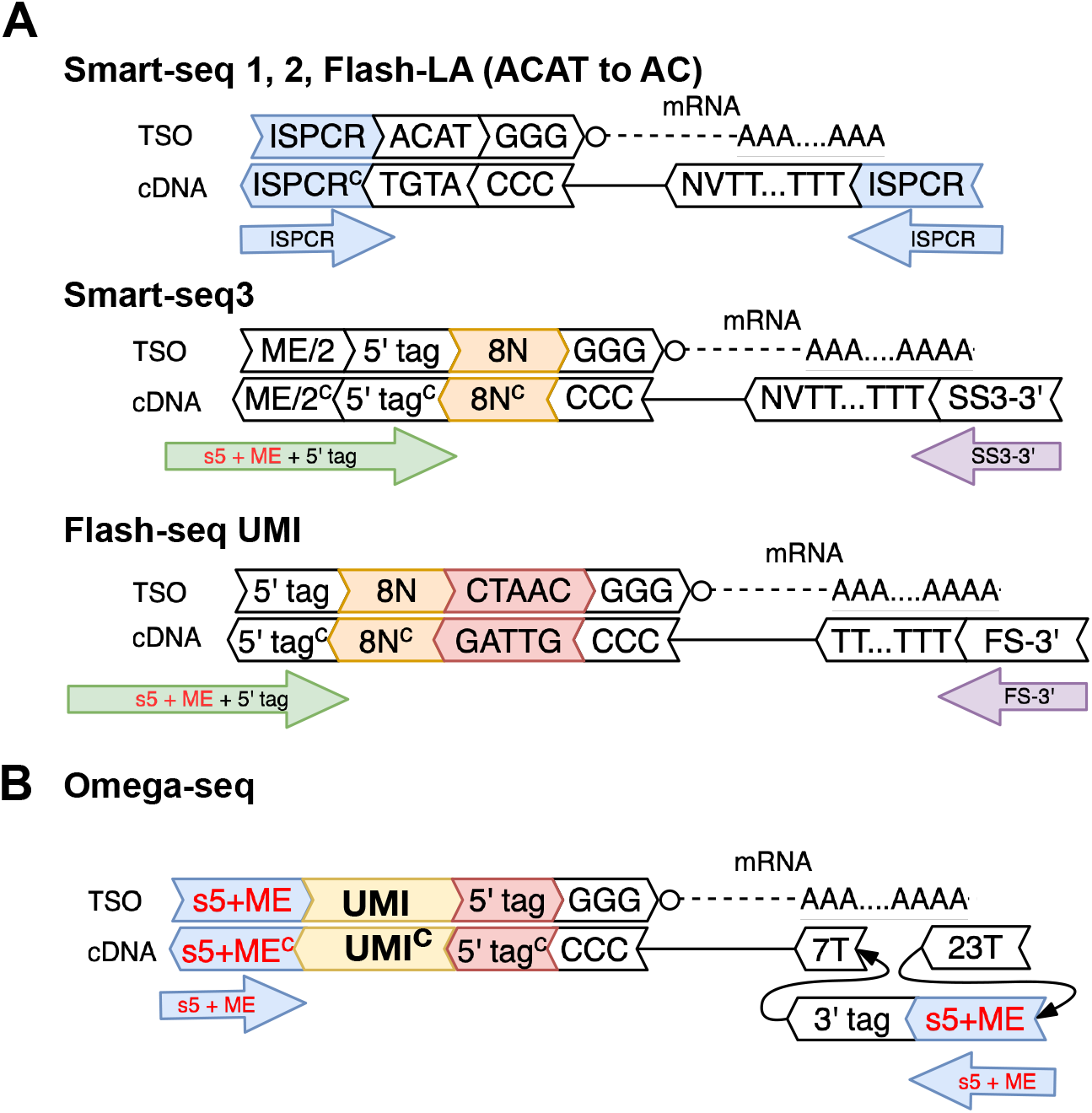
**A**: Schematic of RT and PCR oligonucleotides used in standard Smart-seq protocols. **B**: Schematic of the optimized Omega-seq oligonucleotide design. The UMI sequence is UNNNUNNNUNN.

**Fig. 2.**
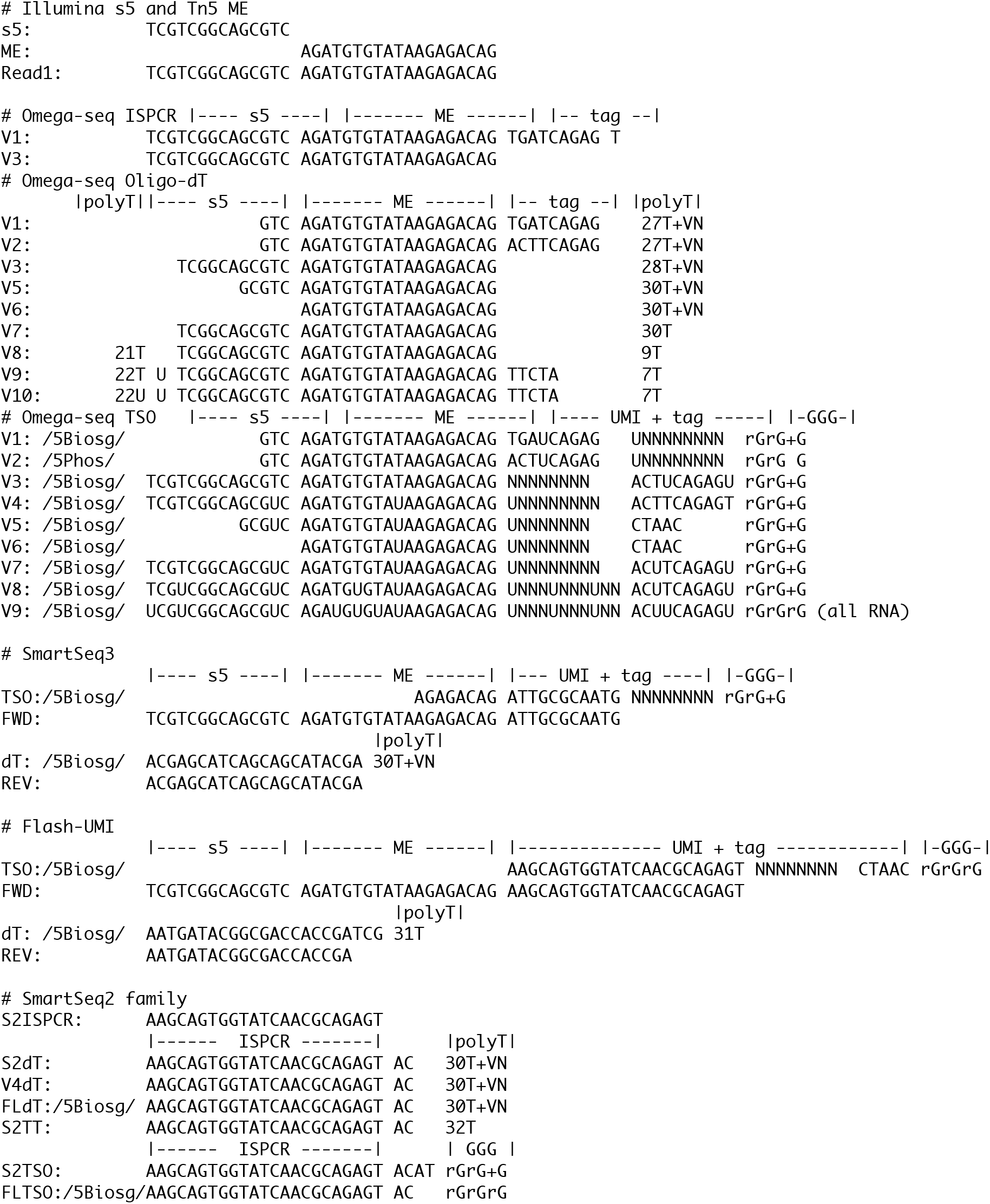
Evolution of Omega-seq oligonucleotide sequences. Various TSO and oligo-dT designs tested during optimization.

**Fig. 3.**
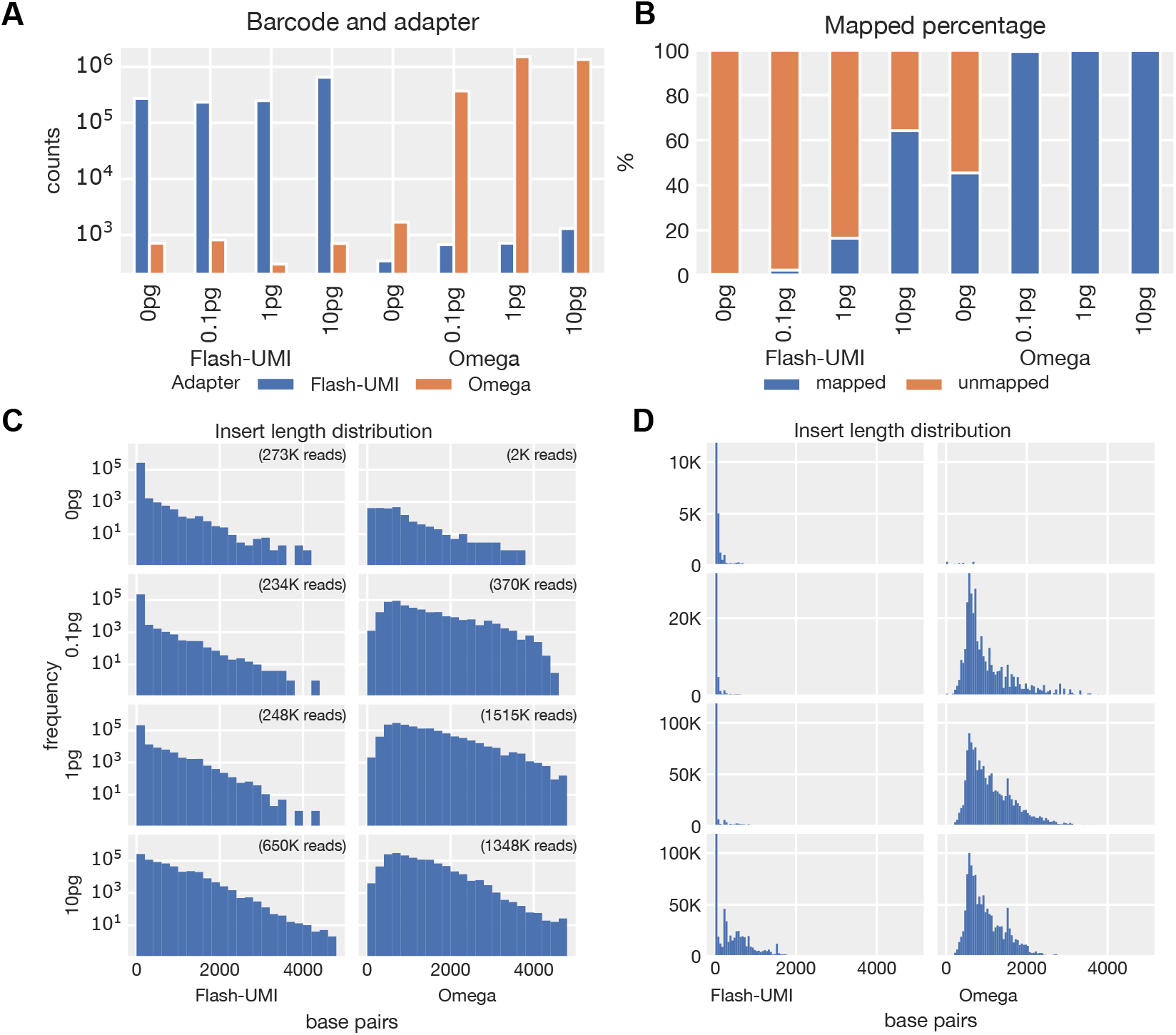
**A**: Number of structured reads according to sample barcodes (x-axis) and adapter type (color). **B**: Percentage of mapped (blue) and unmapped (orange) reads. **C**: Read insert length distribution for FLASH-seq-UMI and Omega-seq Nanopore data in log y-scale. **D**: Same as C but in linear y scale. (Note the first bins of the FLASH-seq-UMI are truncated.)

**Fig. 4.**
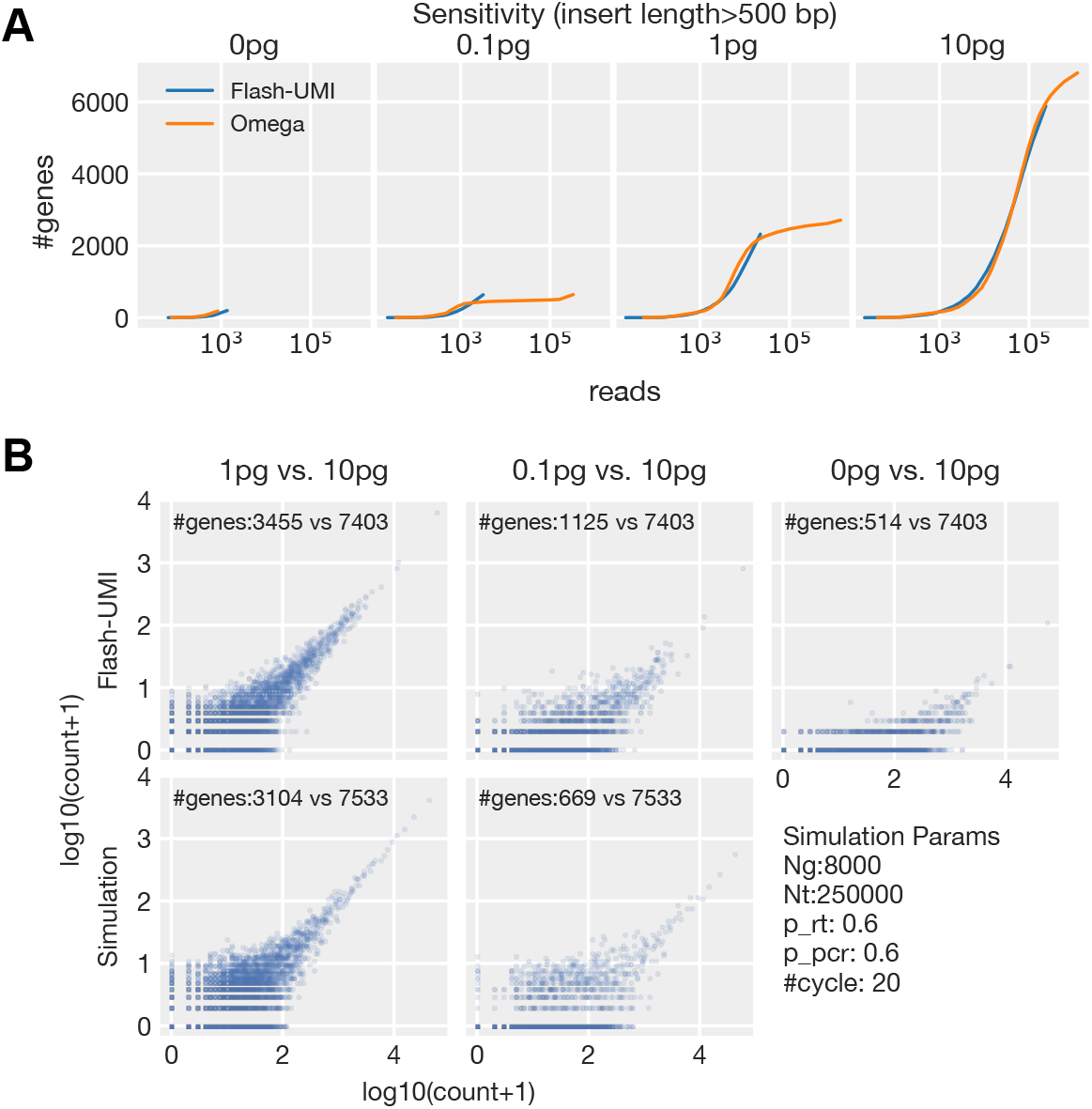
**A**: Similar to Fig. 3E but only reads longer than 500 bp are used. **B**: Similar to Fig. 3F but for FLASH-seq-UMI Nanopore data. The simulation parameters are the same as in Fig. 3F, except for sequencing depth.

**Fig. 5.**
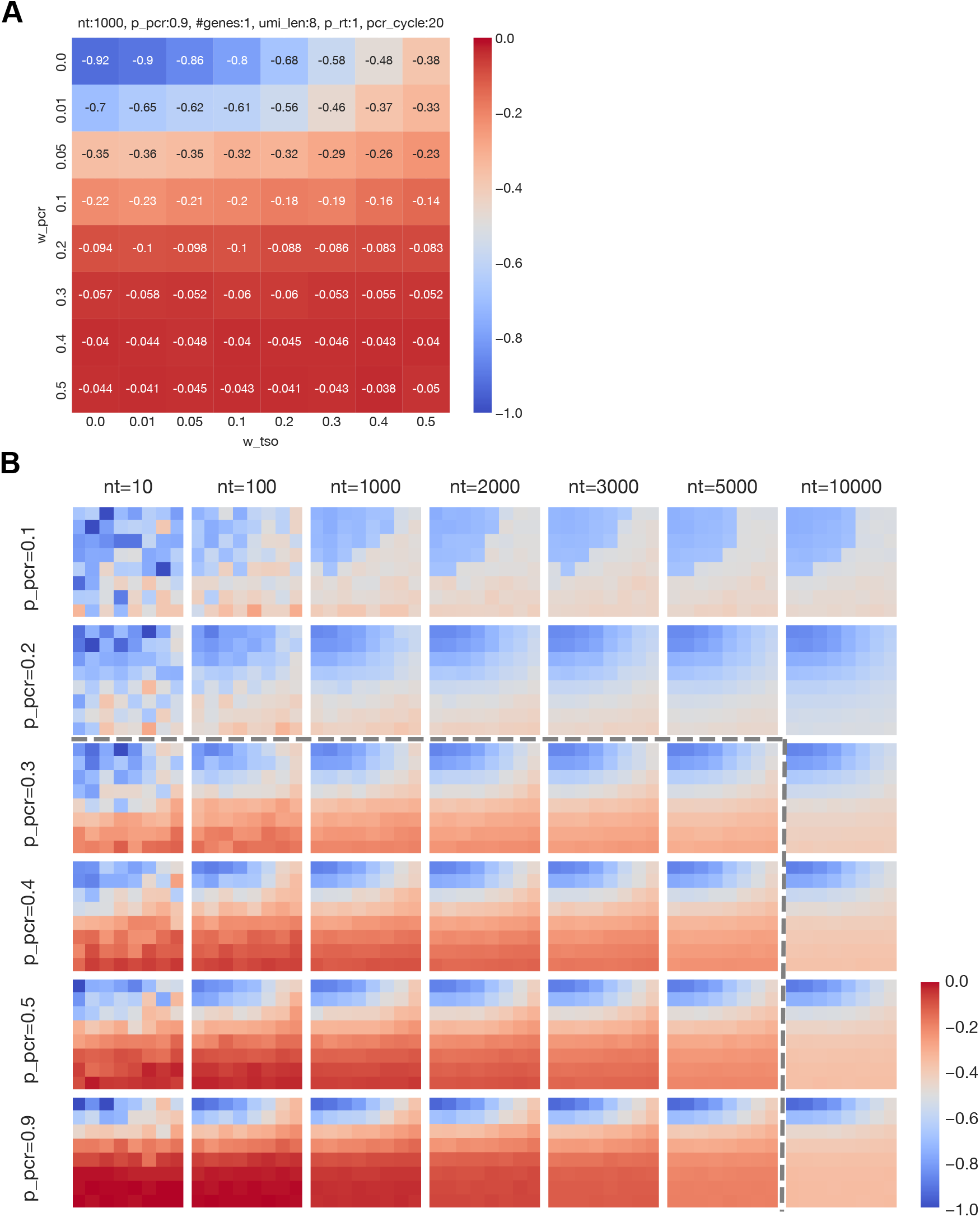
**A**: Simulation of phantom UMI generation across various leak parameters for a single transcript (copy number = 1000) with a baseline PCR efficiency of 0.9. The resulting logQ-slopes are displayed as a heatmap. Notably, the logQ-slope is significantly more sensitive to w_pcr (the binding rate of the TSO to amplicons possessing the proper PCR primer) than to w_tso (the binding rate of the TSO to amplicons already possessing a terminal TSO). This differential sensitivity arises because a positive w_pcr opens the main amplification branch as a continuous source of phantom UMIs, whereas w tso strictly controls the amplification efficiency within the minor, artifactual branch. **B**: Similar simulation to A, but varying the copy number of a single transcript species (*n*_*t*_) and PCR efficiency (p_pcr). At extremely low PCR efficiencies (p_pcr=0.1), the library undergoes only a 5- to 7-fold amplification even after 20 cycles, which roughly translates to the equivalent of 2 to 3 cycles in a high-efficiency PCR. Consequently, phantom UMI generation lacks the exponential depth required to form a heavy-tailed distribution. Conversely, at high transcript copy numbers (e.g., *n*_*t*_ = 10000), the available UMI barcode space is rapidly exhausted; the resulting UMI collisions artificially flatten the distribution, thereby dampening the logQ-slope signal. At extremely low copy number (*n*_*t*_ = 10), the logQ-slope becomes noisy because of the stochastic nature of low-input PCR. However, overall behavior of logQ-slope remains robust and consistent across a wide range of parameter values, even at these extremes.

**Fig. 6.**
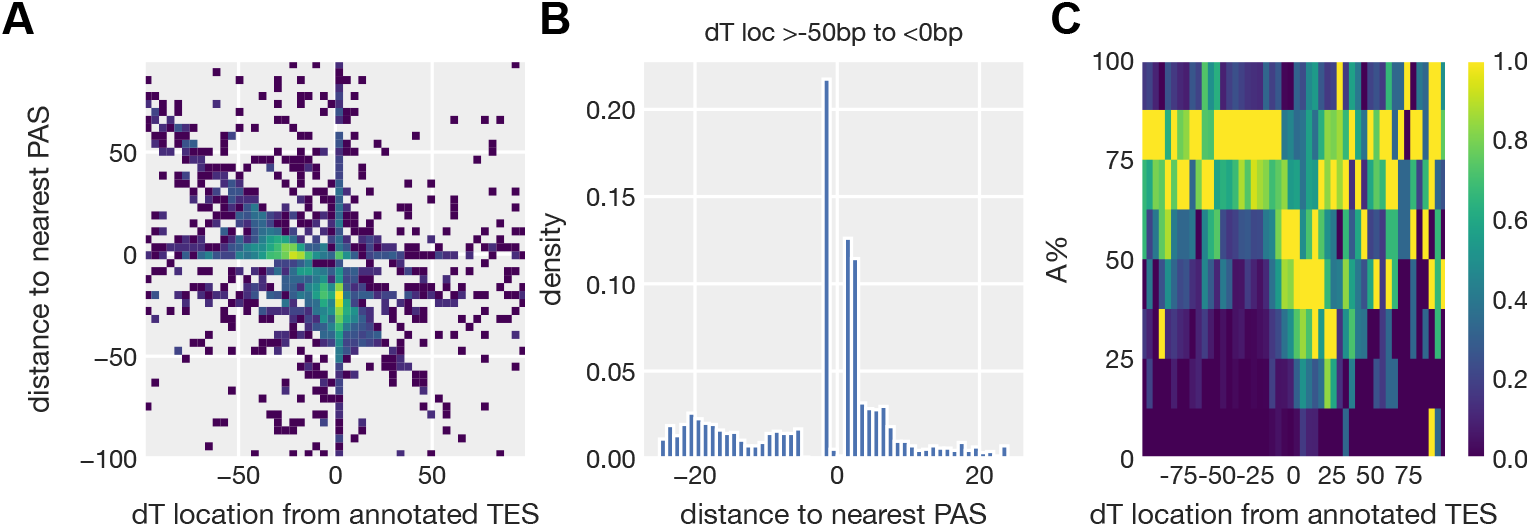
**A**: Joint distribution of dT read position relative to the annotated TES (x-axis, dT − TES) and of the nearest upstream polyadenylation signal relative to the dT read (y-axis, PAS − dT; AATAAA). The PAS typically lies 10–30 bp upstream of the TES, so reads priming at the TES appear near (0, − 25) and reads priming at the PAS near ( − 25, 0). The slope − 1 cluster, along which the PAS–TES spacing is constant, indicates that the nearest PAS is the one associated with that TES. **B**: The dT reads located within 50 bp upstream of annotated TES are enriched near PAS. **C**: Percentage of adenine residues in a 10 bp genomic window downstream of each dT-read position. High adenine content (*>* 75%) indicates internal priming at genomic A-rich tracts.

**Fig. 7.**
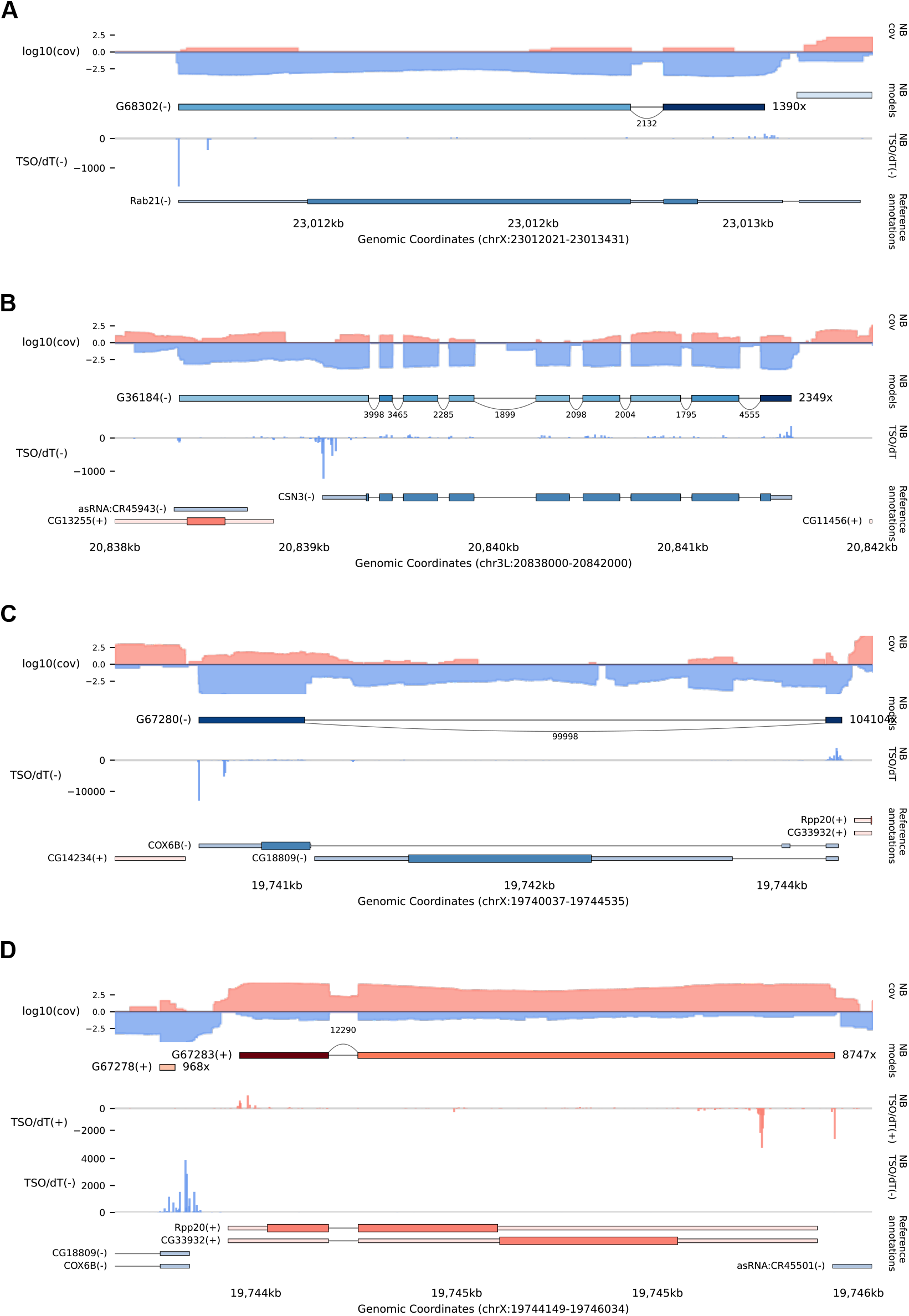
Additional examples of *de novo* reconstruction of gene models, utilizing TSO/dT reads.

**Fig. 8.**
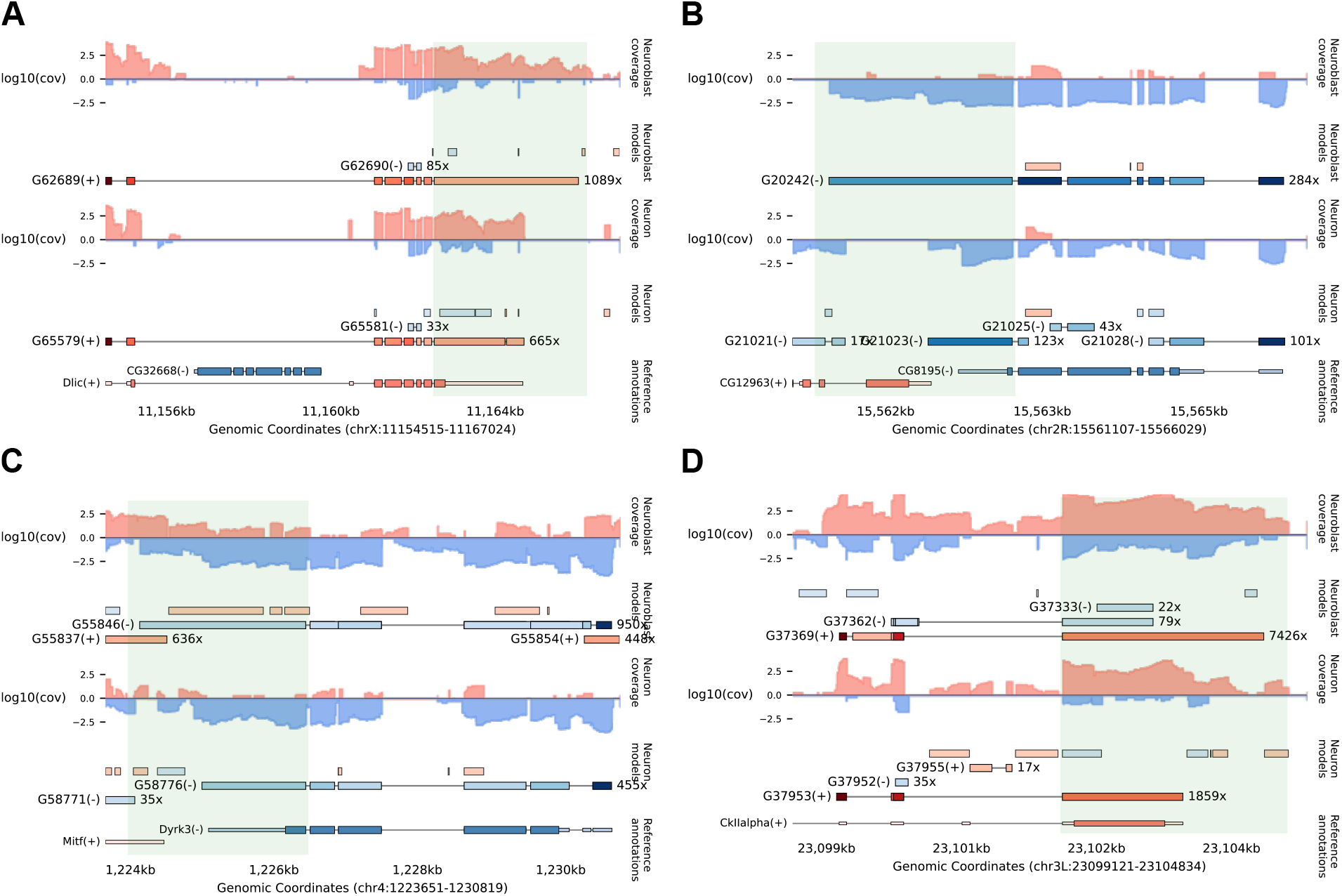
Additional examples of longer unannotated 3′ UTRs, differentially expressed between *Drosophila* neuroblasts and neurons.

**Fig. 9.**
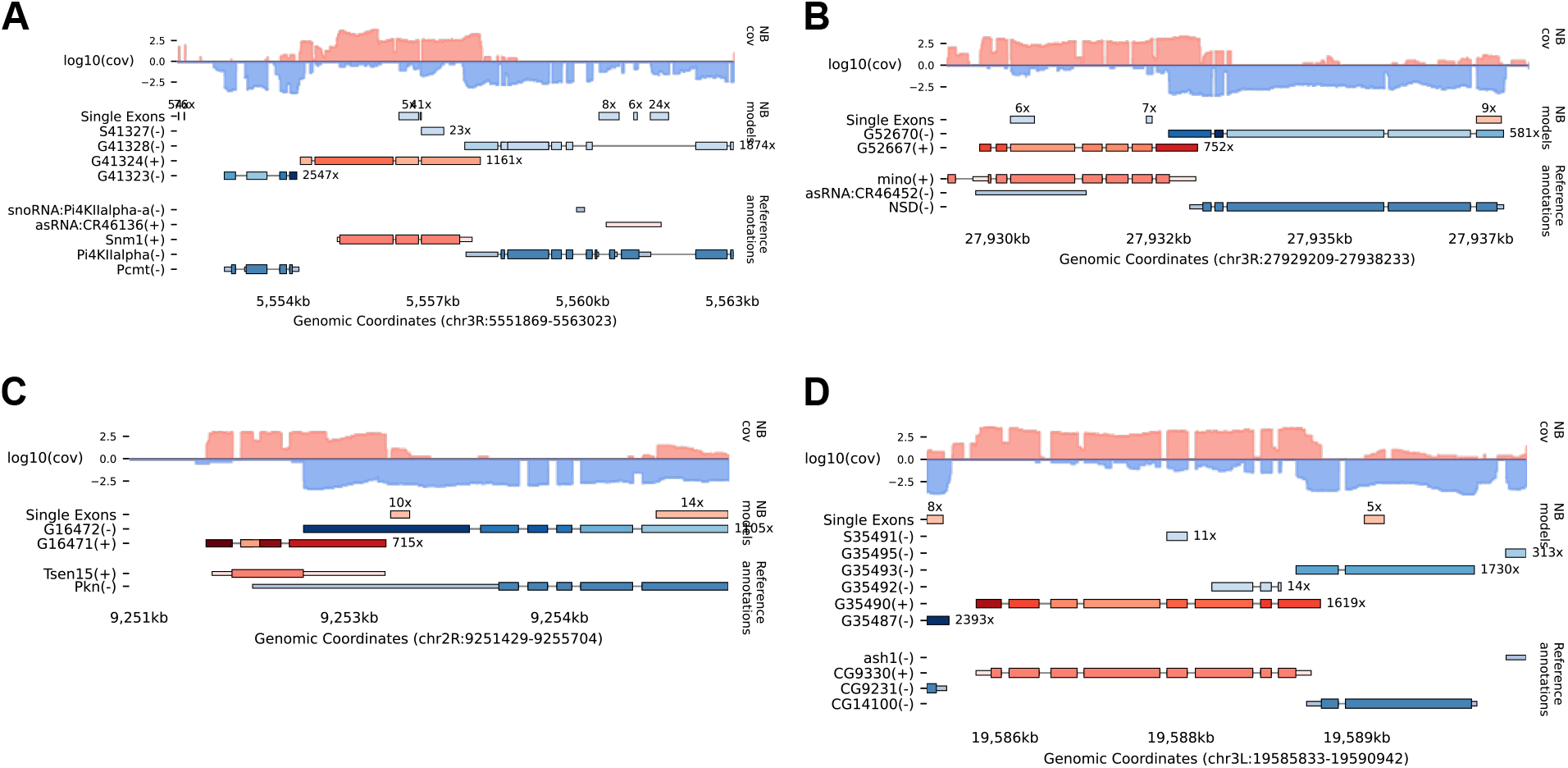
Additional examples of overlapping converging 3′ UTRs.

**Fig. 10.**
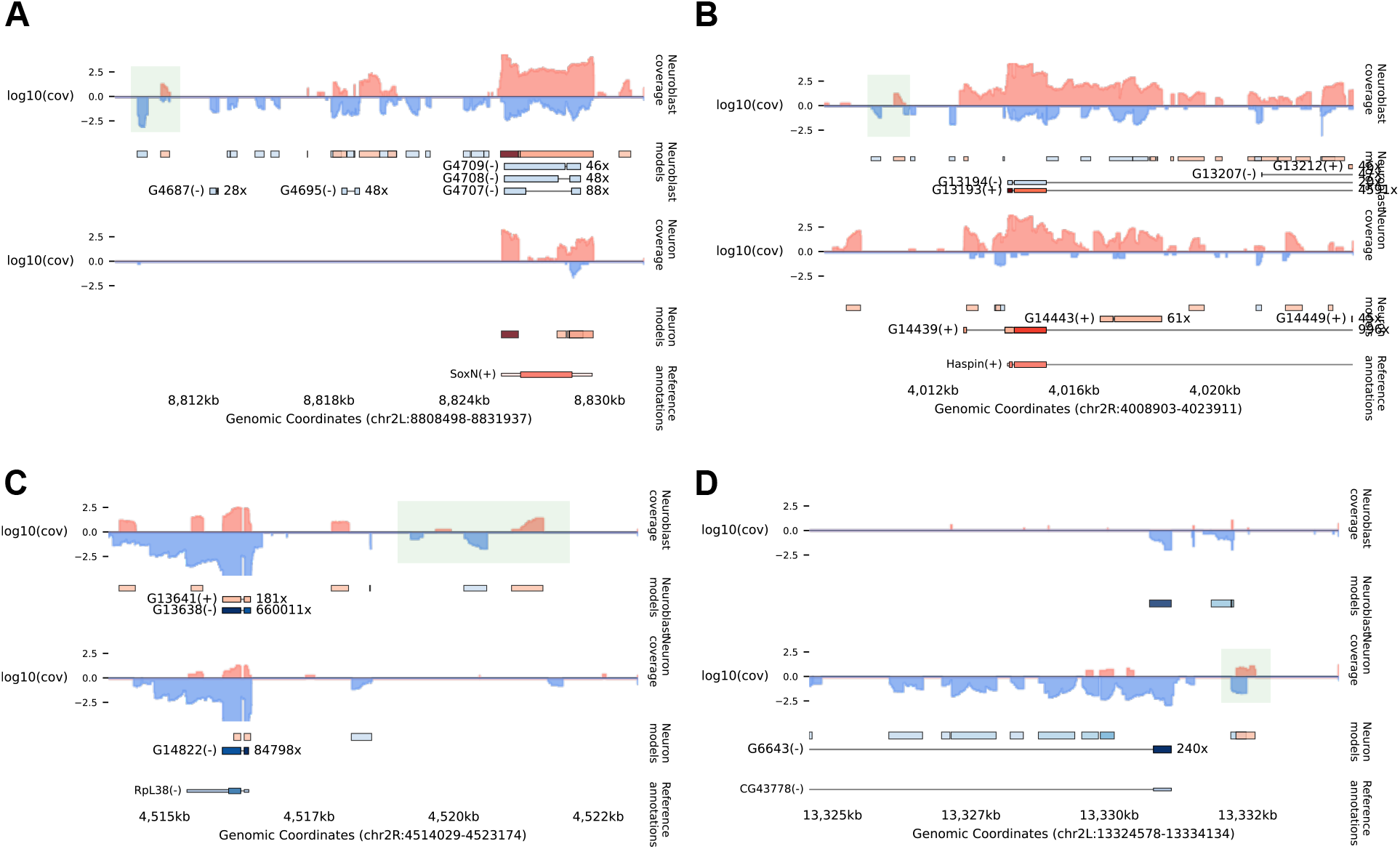
Additional examples of candidate eRNAs.

**Fig. 11.**
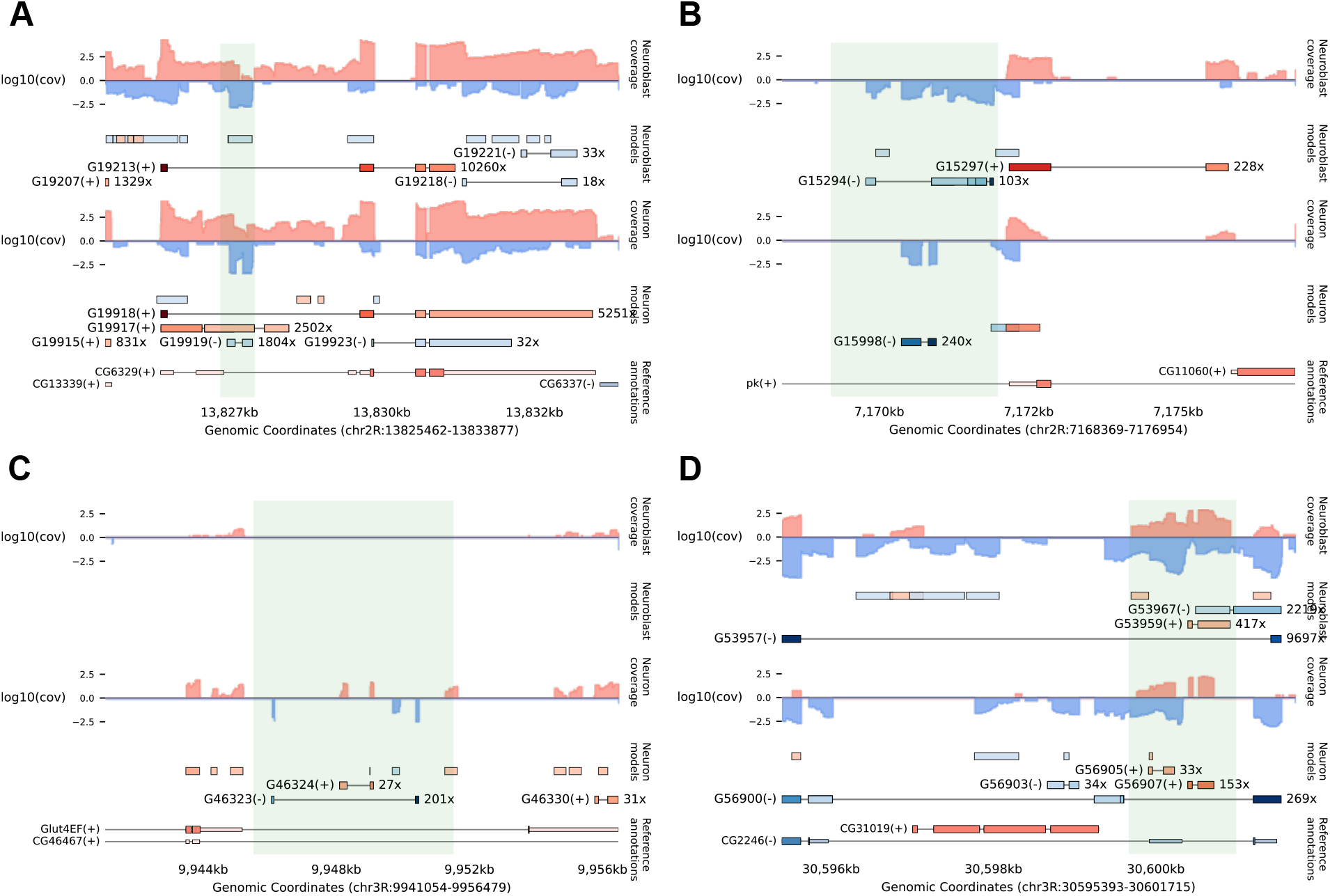
Examples of unannotated antisense transcripts within introns of annotated genes.

**Fig. 12.**
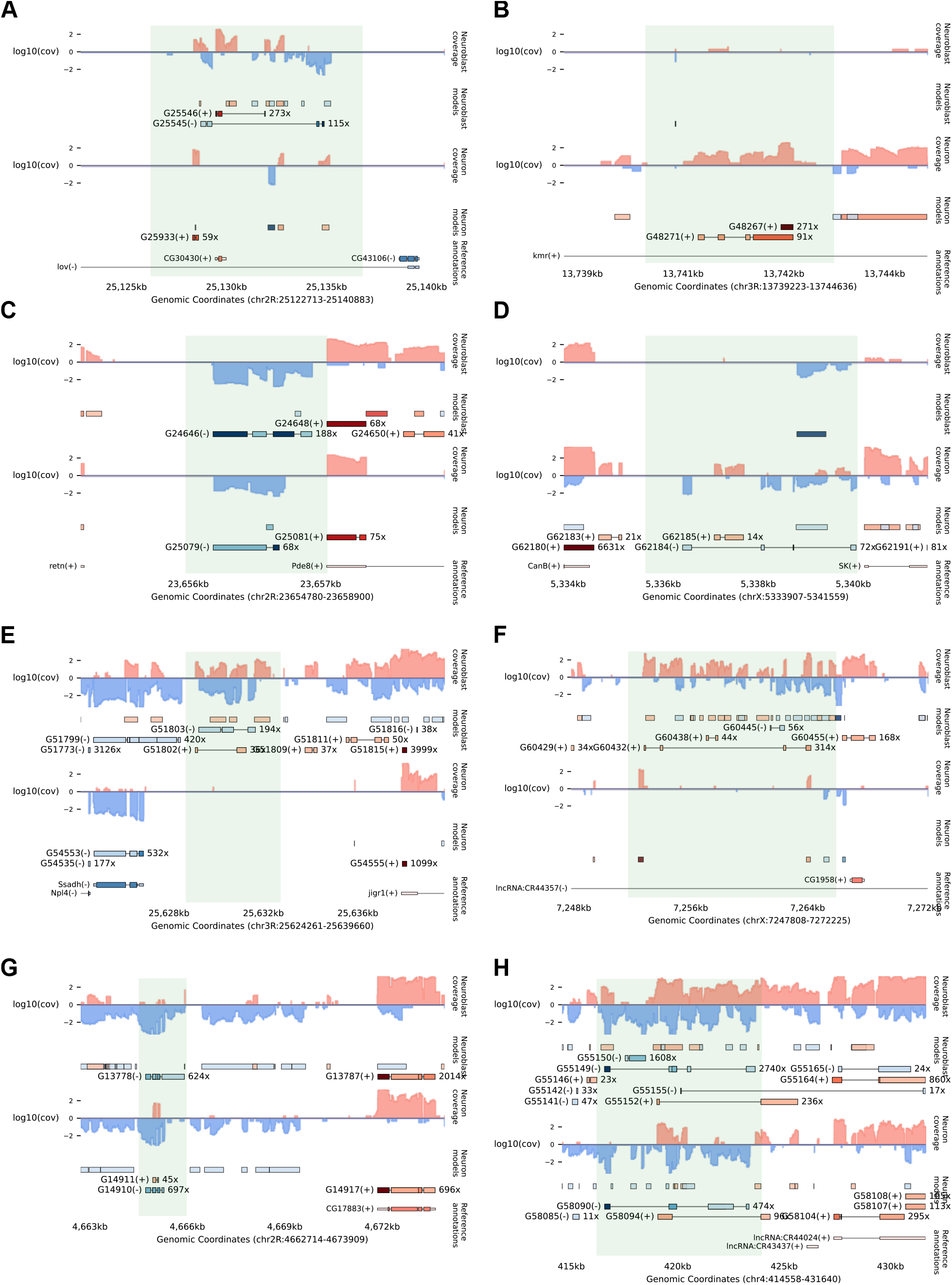
Additional examples of candidate novel genes containing more than three exons.

